# Dynein-Powered Cell Locomotion Guides Metastasis of Breast Cancer

**DOI:** 10.1101/2023.04.04.535605

**Authors:** Yerbol Tagay, Sina Kheirabadi, Zaman Ataie, Rakesh K. Singh, Olivia Prince, Ashley Nguyen, Alexander S. Zhovmer, Xuefei Ma, Amir Sheikhi, Denis Tsygankov, Erdem D. Tabdanov

## Abstract

Metastasis is a principal cause of death in cancer patients, which remains an unresolved fundamental and clinical problem. Conventionally, metastatic dissemination is linked to the actomyosin-driven cell locomotion. However, locomotion of cancer cells often does not strictly line up with the measured actomyosin forces. Here, we identify a complementary mechanism of metastatic locomotion powered by the dynein-generated forces. These forces that arise within a non-stretchable microtubule network drive persistent contact guidance of migrating cancer cells along the biomimetic collagen fibers. We also show that dynein-powered locomotion becomes indispensable during invasive 3D migration within a tissue-like luminal network between spatially confining hydrogel microspheres. Our results indicate that the complementary contractile system of dynein motors and microtubules is always necessary and in certain instances completely sufficient for dissemination of metastatic breast cancer cells. These findings advance fundamental understanding of cell locomotion mechanisms and expand the spectrum of clinical targets against metastasis.

## INTRODUCTION

Metastasis is a process of cancer cell migratory exit from the primary tumor, followed by their invasion into the healthy tissues to establish secondary tumors, which is the leading cause of mortality among cancer patients. For cancer cells, migration in solid tissues represents confined locomotion within and along the available intercellular luminal or interstitial spaces ^1^. Similarly, confined migration in three-dimensional (3D) collagen-rich matrices at the tumor-stroma interfaces occurs *via* paths of least resistance and along the anisotropically aligned collagen fibers ^2^, the phenomenon known as ‘contact guidance’.

Most metastatic cells display a reduced actomyosin contractility in confinement, followed by cell ‘fluidization’ ^3–8^. The question arises on the source of alternative mechanical forces (*i.e.*, non-actomyosin motors) that could facilitate the locomotion of cancer cells in confinement during their breakout from the primary tumor ^9–11^ and infiltration through the blood vessel walls into the healthy tissues ^12^. Answering this question requires a more detailed, higher-order mechanobiological model of cancer cell motility in various conditions, including reduced actomyosin contractility.

The growing body of evidence suggests that microtubules and microtubule-associated motors ^13–15^ are directly involved in the locomotion of cancer cells. For example, migration of mesenchymal cancer cells on collagen-coated surfaces depends on the activity of cytoplasmic dynein motors upon reduction of the actomyosin contractility ^16^, with the latter displaying the instances of interference with the dynein-dependent cell locomotion. Similar observations are reported for cancer cells within the fibrous 3D collagen matrices, where mesenchymal-type cancer cells develop dendritic-like protrusions along the collagen fibers in a dynein-dependent manner ^17^. The pharmacological targeting of dyneins and kinesin-1 that shifts their mechanically antagonistic balance in cancer cells interferes with the organization of microtubules, cell shape, and contact guidance, *i.e.,* their spatial alignment along both topographic and flat anisotropic adhesion cues ^14^.

These findings highlight the importance of the microtubules and their motors for the cell’s ability to sense and interact with its environments. Mechanistically, microtubules are an important cell-scaffolding structure that enables cell alignment to the nano-textures, *i.e.*, *via* direct mechanical and steric interactions of microtubules with the 3D features of the microenvironment ^13^. Moreover, stabilized (*i.e.*, detyrosinated) microtubules feature structural scaffolding and strut-like mechanical function in the cyclically contracting myofibrils within cardiac myocytes ^18^. These observations indicate that the function of microtubules and microtubule-associated motors spans beyond their traditionally accepted role, such as conventional intracellular trafficking, signaling, and cell division.

Here, we aim at uncovering the role of dynein motors in preserving cancer cell motility and contact guidance upon suppressing conventional actomyosin contractility. Specifically, we demonstrate that non-muscle myosin II- and dynein-driven cell contractilities coexist, and each can maintain an effective cancer cell migration along the 2D biomimetic collagen ‘fibers’ network. We also show that cancer cells require the simultaneous activity of dynein and non-muscle myosin II motors during confined 3D migration between densely packed gelatin-based hydrogel microparticles (microgels) within granular hydrogel scaffolds (GHS), highlighting, for the first time, a fundamental mechanobiological difference between confined and unconfined modes of metastasis.

## RESULTS

### Biomimetic substitute of tumor-associated collagen signatures facilitates metastasis studies

The microtubule motors, such as dynein and its cofactors (*e.g.*, dynactin), are robustly linked to cancer progression ^19–24^. For instance, dysregulation of dynactin’s subunit 5 **(Figure 1a**, *left*) or dynein’s heavy chain 1 **(Figure 1a**, *right*) expression is an established marker of a substantially lower survival rate of breast cancer patients ^25^. The upregulation of dynactin subunit 2 (DCTN2) and subunit 5 (DCTN5) expression predicts the metastatic aggressiveness of breast cancer in patients **(Figure 1b)** ^26^. The progression of breast cancer metastatic aggressiveness (*i.e.*, cancer stage progression) also indicates an incremental correlation between such progression and the upregulation of dynactin subunit 2 (DCTN2) and dynein heavy chain 1 (DYNC1H1) **(Figure 1c)** ^26^. Thus, a timely question arises on the specific role of the microtubules (MT) and MT-associated motors in metastasis.

**Figure 1.**
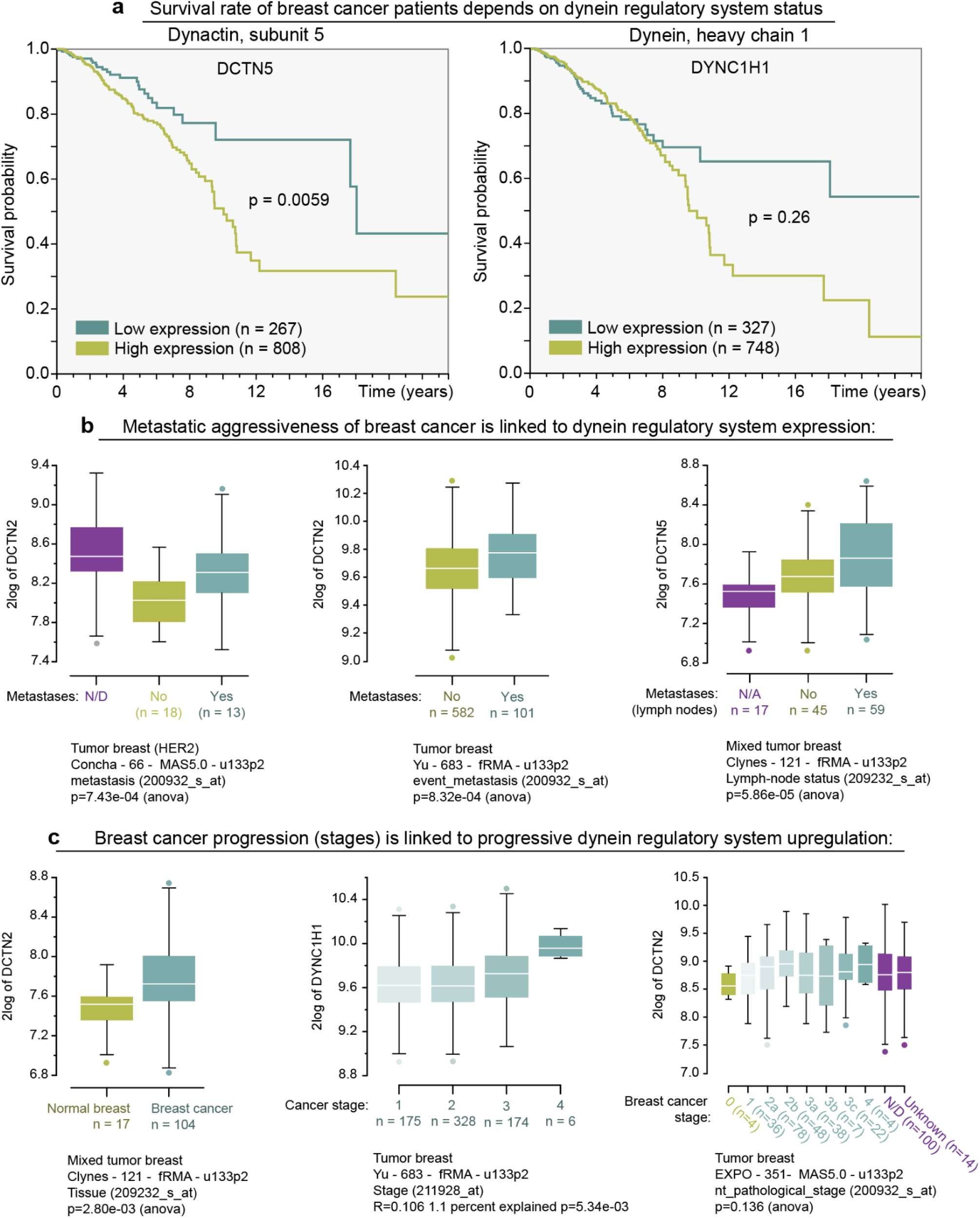
Upregulation of dynein and its regulatory cofactor dynactin’s subunits are linked to the higher mortality rate and metastatic aggressiveness in the breast cancer patients. **(a)** Dynactin’s subunit 5 (DCTN2, *left*) and cytoplasmic dynein’s heavy chain 1 (DYNC1H1, *right*) upregulation indicates a lower survival rate of the breast cancer patients. **(b)** Metastatic aggressiveness of breast cancer is linked to overexpression of dynactin subunits 2 (DCTN2, *left and center*) and 5 (DCTN5, *right*). **(c)** Breast cancer stage progression is linked to the progressive dynein regulatory system upregulation. The binary difference in DCTN2 expression in normal breast *vs.* breast cancer tissues (*left*); progressive increase of the dynein heavy chain 1 overexpression across first (non-metastatic) through fourth (metastatic) breast cancer stages (DYNC1H1, *center*); progressive upregulation of the dynactin subunit 2 across various stages of breast cancer stages (DCTN2, *right*).

Since cancer cells often migrate and exit the primary tumor throughout and along the fibrous collagen-rich extracellular matrices (*i.e.*, ECM), the biomimetic substitutes of ECM, such as 3D collagen gels, became major models for metastasis studies ^9, 15, 27–33^. Nevertheless, the structural and mechanical complexity of 3D collagen gels effectively prohibits detailed mechanobiological studies of the motile behavior of cancer cells in ECM ^34, 35^. A question remains whether actomyosin-driven adhesion and contractility alone are sufficient to coordinate, integrate, and provide a coherent ^36, 37^ and persistent ^38^ cancer cell migration within structurally anisotropic and noisy 3D collagen matrices ^39^.

Therefore, to examine the role of actomyosin and microtubule motors during metastasis, we study MDA-MB-231 cells’ migratory behavior on the soft (shear modulus G′=8.6 kPa) and rigid (G′=55 kPa) 2D biomimetic substrates, which consist of flat 1 µm-wide parallel collagen type-1 lanes, *i.e.*, artificial collagen ‘fibers’ that mimic naturally occurring collagen architectures ^2, 29, 40^. We chose the pitching distance between each set of the parallel collagen lane at 15 µm, as it provides a sufficient spatial sparsity of the adhesion cues to prevent the loss cell’s sensitivity to the guidance effects of the collagen architecture ^13, 15, 41^. Moreover, we arrayed two sets of the described collagen lanes into the rhomboid grids, crisscrossed at 22° angle **(Figure 2a-c)**, to mimic anisotropically oriented yet structurally noisy metastasis-driving tumor-associated collagen signatures (*i.e.*, TACS-3 ^2^), routinely observed at the tumor-stroma interface ^2, 10^.

**Figure 2.**
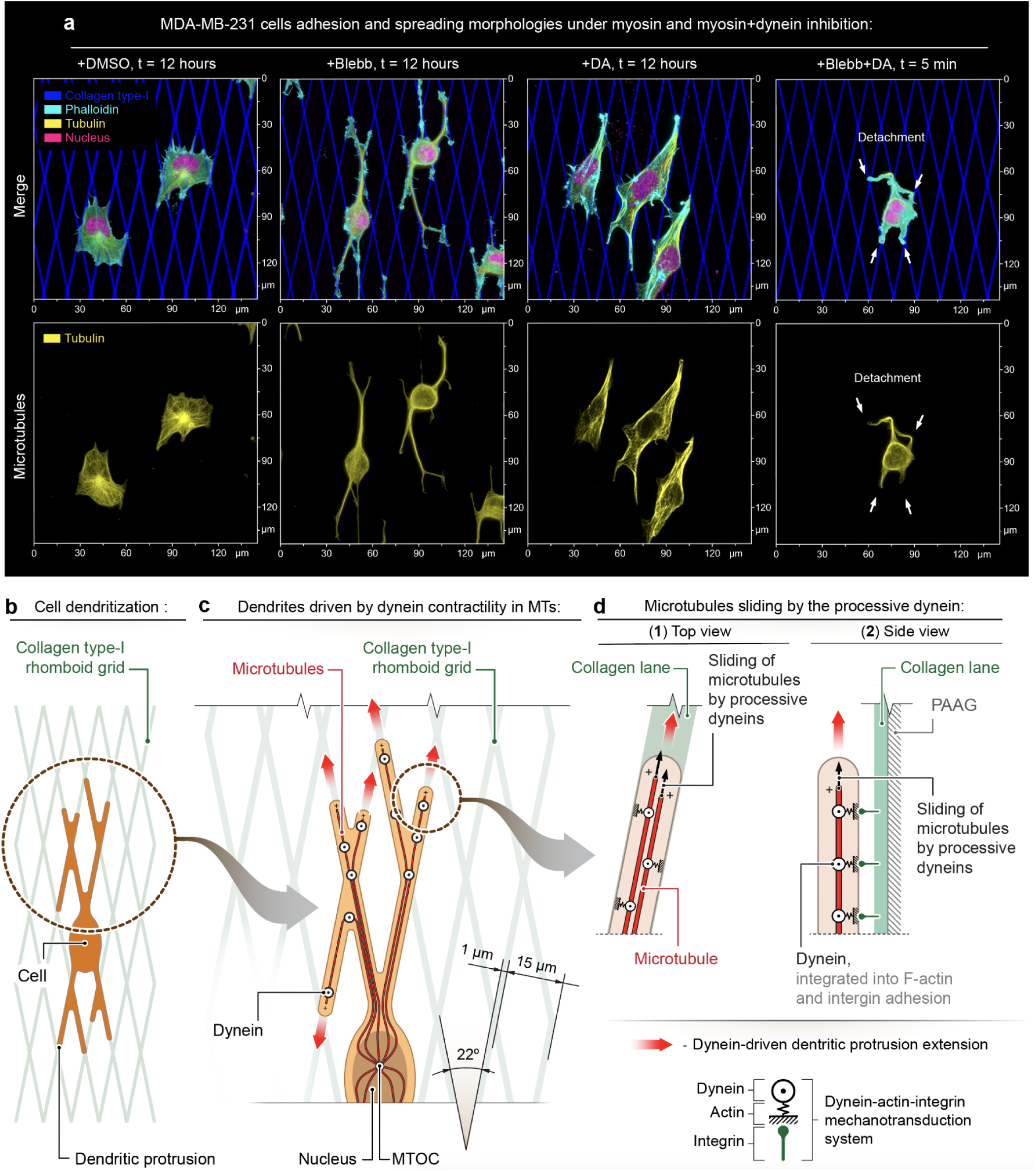
Adhesion and spreading of metastatic MDA-MB-231 cells along the collagen type-1 patterns depend on actomyosin and microtubules-dynein alternative contractility systems. **(a)** MDA-MB-231 morphological reorganization in response to adhesion and spreading along the anisotropic collagen ‘fibers’ (*i.e.*, rhomboid grid)in the control conditions (+DMSO), during suppression of actomyosin contractility with Blebbistatin (+Blebb), during Dynarrestin-induced dynein-microtubules dissociation (+DA), and upon dynein-MT dissociation with Dynarrestin during a prolonged suppression of actomyosin contractility with Blebbistatin (+Blebb+DA). *Note the dendritic architecture of cells during Blebbistatin-induced suppression of actomyosin contractility (+Blebb) and a high level of structural conformity of cell protrusions to the underlying collagen grid. Structural conformity indicates protrusions’ adhesion to the collagen grid along their entire lengths.* **(b)** Schematic view of the dendritized MDA-MB-231 cell on the collagen rhomboid grid during actomyosin contractility suppression. **(c-d)** Suggested schematic representation of the collagen-guided elongation of dendritic-like protrusions along the rhomboid grid. *Note that microtubules structurally scaffold and support the microtubules.* **(c)** Elongation of dendritic-like protrusions (*red arrows*) along the collagen lanes. *Note the structural characteristics of the collagen rhomboid grid: two sets of 1 µm-wide 15 µm-pitched parallel collagen lanes, criss-crossed into a rhomboid grid under 22° angle*. **(d)** Dendritic-like protrusions slide along the collagen ‘fibers’ via the dynein motors activity that slide the dendritic protrusions’ structural core (i.e., microtubules). The mechanical coupling between the microtubules to the guiding collagen lanes is enabled via dynein motors, embedded into the F-actin cytoskeleton that in turn is coupled to the ECM via the adhesion apparatus (e.g., integrin system).

### Low actomyosin contractility reveals that cell adhesion and spreading depend on dyneins

We test MDA-MB-231 cells adhesion and spreading in control conditions (empty vehicle, **Figure 2a**, **+DMSO)**, low actomyosin contractility (+Blebbistatin, 12 hours, **Figure 2a**, **+Blebb)**, low dynein activity (+Dynarrestin ^14, 42^, 12 hours, **Figure 2a**, **+DA)**, and after subsequent 12-hour-long blebbistatin treatment, followed by 5 min-long co-treatment with dynarrestin (DA) **(Figure 2a**, **+Blebb+DA)**. Inhibition of principal locomotion machinery, *i.e.*, actomyosin contractility ^43^, by targeting non-muscle myosin II with blebbistatin ^44, 45^ does not suppress cell-grid adhesion. Compared with the compact polygonally-shaped control cells **(Figure 2a**, **+DMSO, Movie 1)**, blebbistatin treatment increases cell spreading and induces the formation of the branched protrusions along the collagen grid **(Figure 2a**, +Blebb, Movie 2).

To test the dynein-driven cell spreading, we use dynarrestin (DA), which is a selective dynein inhibitor that prevents dynein-MT interactions ^42^, featuring a higher inhibiting efficiency than the ciliobrevin family ^46^. We dismiss Dynapyrazole A as a dynein inhibitor because, unlike DA, it blocks dynein’s processive activity along the MTs but does not prevent dynein-MT interactions, rendering dyneins into the passive MT crosslinkers ^47^. Inhibition of dyneins in the presence of intact actomyosin contractility does not detectably affect MDA-MB-231 cell’s ability to adhere and spread along the collagen grids **(Figure 2a**, **+DA)**. Cells preserve the polygonal morphology throughout the entire course of a 12-hour-long DA treatment, similar to the control cells **(Figure 2a**, **+DMSO** *vs.* **+DA)**. On the contrary, dynein suppression in cells with a low actomyosin contractility (blebbistatin-pretreated, 12 hours) abruptly dismantles the blebbistatin-induced dendritic protrusions **(Figure 2a**, **+Blebb+DA)**, followed by cells detachment from the collagen grids **(Movie 3)**. Thus, in the absence of substantial actomyosin contractility, dynein activity becomes indispensable for maintaining stable dendritic protrusions.

Our results indicate that either actomyosin or dynein motors are able to provide cell spreading while disabling both motor systems results in a complete loss of cell adhesion. Based on these observations, we suggest a model of locomotion powered by dynein-driven contractility with the microtubules, which provides cell adhesion, spreading, and motility **(Figure 2b-d)**. In this model, cells with decreased actomyosin contractility **(Figure 2b)** utilize the mechanically loaded microtubule cables to transmit dynein-generated forces along dendritic protrusions **(Figure 2c)**, rendering microtubules into the mechanically and structurally active components of the cytoskeleton ^13, 14, 18, 48^. Since microtubules mechanically integrate into the F-actin network *via* the dynactin complex, an F-actin-mimicking dynein cofactor ^49–52^, we also suggest that branched F-actin cytoskeleton may serve as an adaptor between microtubules and integrin-based cell adhesion machinery for the microtubules+dynein-driven cell adhesion, spreading, and locomotion **(Figure 2d)**. Indeed, dynactin is a nucleation seed for Arp2/3 complex assembly ^51^ that, in turn, nucleates the branched F-actin network. Thus, the dynein-dynactin complex acts as a supporting system for the low-contractility branched F-actin network that grows around the microtubules in dendritic protrusions, while inhibition of Arp2/3 in these cells with CK666 results in protrusion failure and cell detachment ^15^.

### Simultaneous inhibition of dynein and actomyosin contractility arrests locomotion of metastatic cells

We examine changes to MDA-MB-231 cell migration **(+DMSO**, ∼12 hours) in response to **(i)** suppression of the actin-myosin **(+Blebbistatin**, ∼12 hours), later combined with inhibition of microtubules-dynein (+Dynarrestin) cytoskeletal motor subsystems **(Figure 3a-c**, **+DMSO→+Blebb→+Blebb+DA, Movie 4)**, or **(ii)** suppression of the dynein activity first (+Dynarrestin, 12 hours) followed by co-inhibition of non-muscle myosin II (+Blebbistatin) **(Figure 3d-e**, **+DMSO→+DA→+DA+Blebb, Movie 5)**.

**Figure 3.**
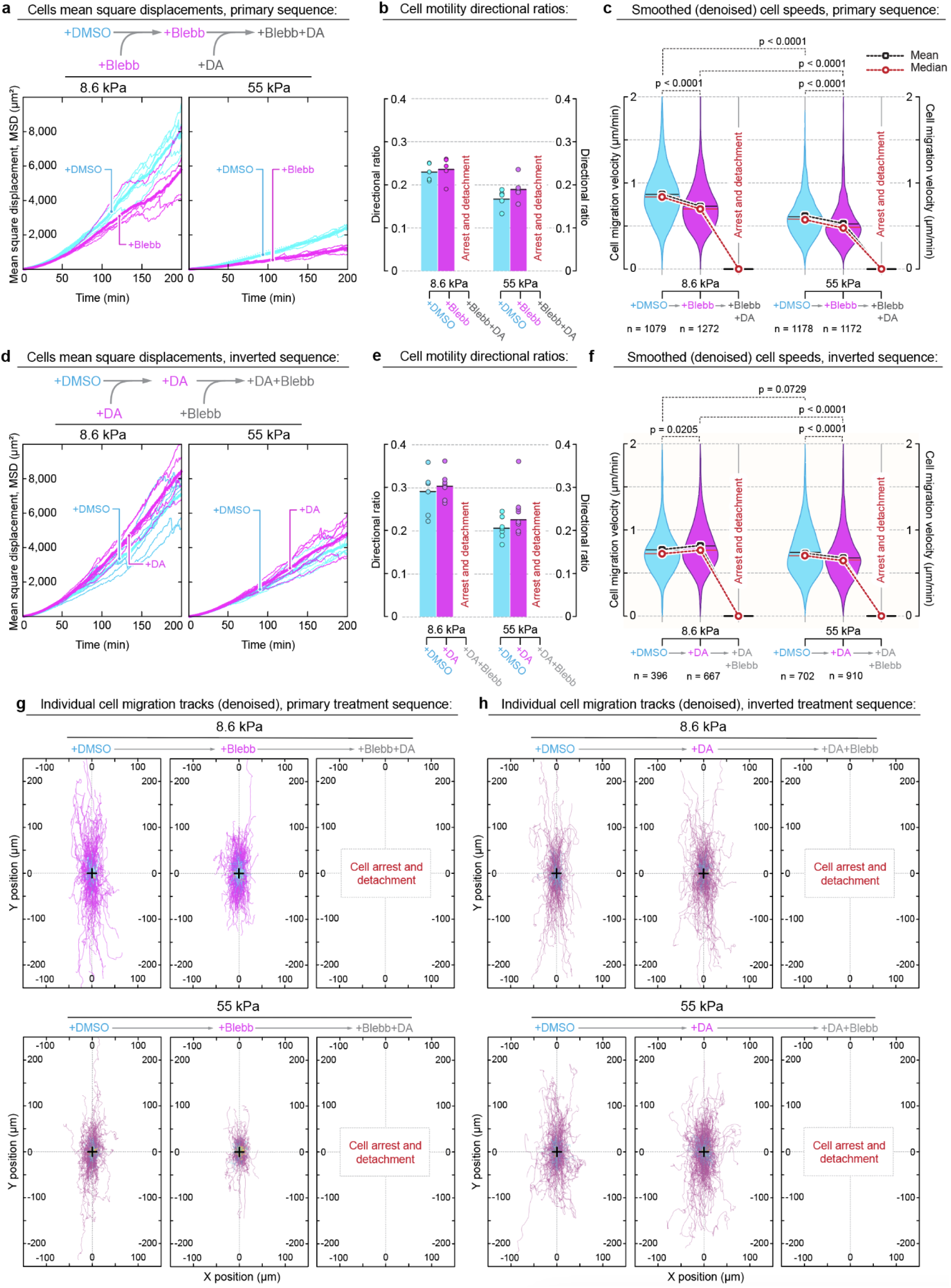
Metastatic MDA-MB-231 cells use parallelized myosin- and dynein-driven systems to power collagen-directed migration and contact guidance. **(a-c)** Live cell migration imaging and analysis are performed sequentially (primary sequence) within the same cell samples and fields of view as follows: *First* - control conditions upon addition of the vehicle (+DMSO, *cyan*); *Second* - during actomyosin contractility inhibition upon addition of Blebbistatin (+Blebb, *magenta*); *Third* - dynein suppression upon addition of Dynarrestin over Blebbistatin (+Blebb+DA, *gray*). *Note that MDA-MB-231 cells in both control (+DMSO) and during low actomyosin contractility (+Blebb) states migrate at longer distances on the soft, collagen ‘fibers’ (G′=8.6 kPa). Note that MDA-MB-231 cells do not display any migratory properties (arrest and detachment) in Blebbistatin+Dynarrestin mixture.* **(a)** Cell mean square displacements (MSD) of MDA-MB-231 cells migrating along the soft (G′=8.6 kPa, *left*) and rigid (G′=55 kPa, *right*) collagen grids. **(b)** Directional ratios of the migrating MDA-MB-231 cells along the collagen grids. **(c)** Smoothed (*i.e.*, denoised) MDA-MB-231 cell migration speeds across all treatment conditions. *Note that suppression of actomyosin contractility with Blebbistatin does not arrest, but only partially slows down migration of MDA-MB-231 cells. Note that MDA-MB-231 cells in control conditions (+DMSO) migrate with a higher speed on the soft (G′=8.6 kPa), than on the rigid (G′=55 kPa) collagen ‘fibers’. On the contrary, upon actomyosin contractility suppression with Blebbistatin (+Blebb) dendritic-like MDA-MB-231 cells (i.e., dynein-dependent) display higher speeds on the rigid (G′=55 kPa) collagen grids.* **(d-f)** Sequential analysis of continuous cell migration across the following conditions (inverted sequence): *First* - control conditions (+DMSO, *cyan*); *Second* - during dynein motor protein activity inhibition upon addition of Dynarrestin (+DA, *magenta*); *Third* - actomyosin contractility suppression upon addition of Blebbistatin atop of Dynarrestin (+DA+Blebb, *gray*), *i.e.*, +DMSO→+DA→+DA+Blebb sequence. **(d)** Mean square displacements of the MDA-MB-231 cells MSD on soft (G′=8.6 kPa, *left*) and rigid (G′=55 kPa, *right*) collagen grids during the course of the sequential ‘inverted’ treatments (*i.e.*, +DMSO→+DA→+DA+Blebb). *Note that dynein inhibition (+DA) does not significantly affect MDA-MB-231 cell displacement from the originating point, compared to the same cells in control conditions (+DMSO).* **(e)** Directional ratios of the migrating MDA-MB-231 cells along the collagen grids across all treatment conditions. **(f)** Denoised speeds of MDA-MB-231 cells across all treatment conditions. *Note that dynein inhibition **(+DA)** does not significantly affect cell migration speeds, compared to the same cells in control conditions **(+DMSO)***. **(g)** Individual smoothed (*i.e.*, denoised) migration tracks of the MDA-MB-231 cells on soft (G′=8.6 kPa, *top*) and rigid (G′=55 kPa, *bottom*) collagen grids across +DMSO→+Blebb→+Blebb+DA conditions. (h) Denoised) migration tracks of the MDA-MB-231 cells on soft (G′=8.6 kPa, *top*) and rigid (G′=55 kPa, *bottom*) collagen grids across +DMSO→+DA→+DA+Blebb conditions. *Note that in both +Blebb+DA and +DA+Blebb MDA-MB-231 cells experience the migration arrest, followed by the cell detachment from soft and rigid grids*.

In the first series of experiments **(i)**, inhibition of actomyosin contractility only partially decreases population-wide effective migratory cell displacement, *i.e.*, mean square displacement (MSD) **(Figure 3a**, **+DMSO** *vs.* **+Blebb, Movie 4)**. The population-wide directional ratio of migrating cells does not change significantly upon addition and incubation in blebbistatin **(Figure 3b**, **+DMSO** *vs.* **+Blebb)**, while the population-wide cell speeds on both soft (G′=8.6 kPa) and rigid (G′=55 kPa) collagen grids only decrease by Δ<10% **(Figure 3c**, **+DMSO** *vs.* **+Blebb)**. The consequent co-inhibition of the dynein with dynarrestin, *i.e.*, in addition to blebbistatin, leads to an abrupt arrest of cell motility **(Figure 3c**, **+Blebb+DA, Movie 3)**, followed by the cell’s dendritic protrusion failure and cell detachment from collagen grids **(Figure 2a**, **+Blebb+DA)**.

In the second series of experiments **(ii)**, suppression of the dynein activity first (+Dynarrestin) also does not significantly affect MDA-MB-231 cell migratory behavior in comparison to the control conditions (+DMSO) on both soft (G′=8.6 kPa) and rigid (G′=55 kPa) collagen grids **(Figure 3d-f**, **+DMSO** *vs.* **+DA, Movie 5)**. Specifically, inhibition of dynein activity on both soft (G′=8.6 kPa) and rigid (G′=55 kPa) collagen grids causes an insignificant increase of the population-wide cell displacement (MSD) **(Figure 3d**, **+DMSO** *vs.* **+DA)**, cell motility directional ratios **(Figure 3e**, **+DMSO** *vs.* **+DA)**, and negligible change of the population-wide cell migration speeds **(Figure 3f**, **+DMSO** *vs.* **+DA)**. The following co-inhibition of the actomyosin contractility with blebbistatin in addition to the preceding ∼12-hour-long dynein suppression (+DA) results in cell arrest **(Figure 3f**, **+DA+Blebb)**. Thus, our results show that separate inhibition of either non-muscle myosin II (+Blebb) or dynein (+DA) motors does not arrest the migration of MDA-MB-231 cells along soft (G′=8.6 kPa) and rigid (G′=55 kPa) collagen grids. However, simultaneous suppression of actomyosin and dynein motors abrogates the migration of cancer cells.

### Isolated myosin- or dynein-driven locomotion supports contact guidance of metastatic cells

To analyze the directional responsiveness of MDA-MB-231 cancer cells to the geometry of the anisotropically oriented collagen grid, *i.e.*, contact guidance ^13, 15, 29, 53^, we evaluate individual tracks of migrating cells **(Figure 3**, **g** and **h)** and the total angle distribution of 1 minute-long cell displacement in respect to the grid’s anisotropy axis **(Figure S1)** for all used conditions.

Our results indicate that the directionality of cell migration is equally responsive to the anisotropy of the collagen contact guidance cues (grids) on soft (G′=8.6 kPa) and rigid (G′=55 kPa) substrates in control conditions **(Figure 3**, **g** and **h, +DMSO, G′=8.6 kPa** *vs.* **55 kPa, Movies 4** and **5)**, during inhibition of the actomyosin contractility **(Figure 3g**, **+Blebb, 8.6 kPa** *vs.* **55 kPa, Movies 4** and **5)**, or during the suppression of the dynein **(Figure 3h**, **+DA, 8.6 kPa** *vs.* **55 kPa, Movies 4** and **5)**.

Similarly, the angle distributions of cell displacements in relation to the grid anisotropy show no significant deflection of the cell migration course from the directionality of both soft (G′=8.6 kPa) and rigid (G′=55 kPa) substrates **(Figure S1, a-b, G′=8.6 kPa** *vs.* **55 kPa)** for all treatment conditions **(Figure S1, a-b, +DMSO, +Blebb**, and **+DA)**. Thus, our data demonstrate that migrating MDA-MB-231 cells can sustain contact guidance *via* myosin- and dynein-driven mechanisms.

### Upregulation of Kinesin-1 activity attenuates dynein-driven locomotion

To further challenge our model of dynein-driven cell motility, we use mechanical antagonism between dynein and kinesin-1 as an orthogonal test to the inhibition of the dyneins in cells with suppressed actomyosin contractility **(Figures 2** and **3)**. For that purpose, we study the migratory response of cancer cells to the kinesin-1 overactivation induced by a small kinesin-1-specific activator, kinesore (KS) ^14, 54^.

The kinesore addition to the blebbistatin-pretreated MDA-MB-231 cells **(Figure 4a-c**, **+DMSO**→**+Blebb)** results in a substantial slow-down of cell locomotion along both soft (G′=8.6 kPa) and rigid (G′=55 kPa) collagen grids **(Figure 4a-c**, **+Blebb**→**+Blebb+KS, G′=8.6 kPa** *vs.* **55 kPa, Movie 6)**. Thus, stimulation of kinesin activity and its mechanical antagonism with dyneins ^14^ indeed interfere with the dynein-driven cell locomotion. Moreover, since kinesore does not inhibit dynein but only over-activates kinesin-1, we do not observe detachment of cells, as opposed to dynarrestin addition to the blebbistatin-pretreated MDA-MB-2312 cells **(Figures 2a** and **3c**, **+Blebb+DA)**.

**Figure 4.**
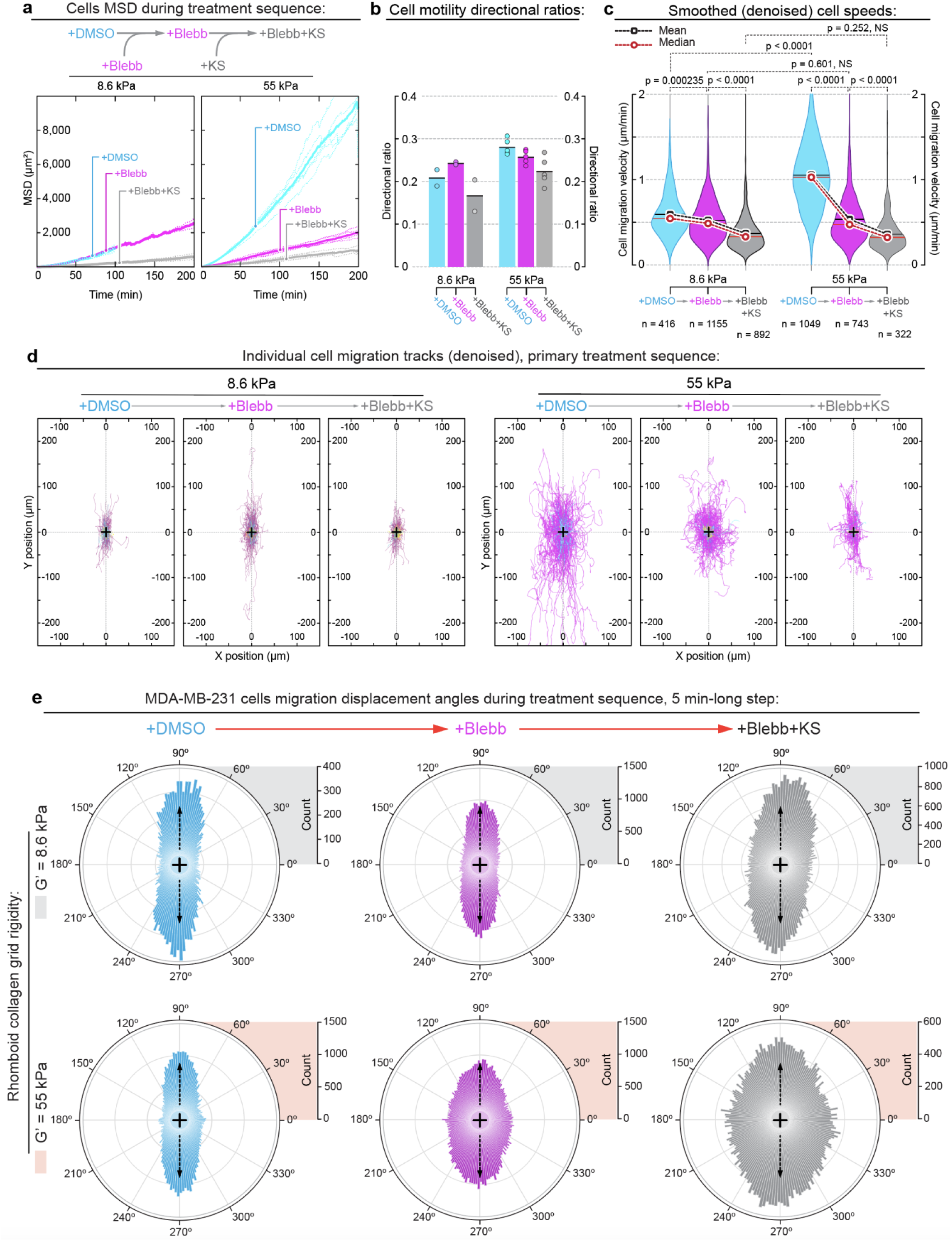
Orthogonal testing of the dynein-driven metastatic locomotion in MDA-MB-231 cells with kinesore. Overactivation of dynein’s mechanical antagonist, kinesin-1 *via* treatment with Kinesore **(+KS)** reduces cell locomotion, but does not significantly affect the contact guidance along the collagen grid. *Note that MDA-MB-231 cells do not completely lose migratory properties and do not detach during combined actomyosin contractility suppression and kinesin-1 overactivation **(+Blebb+KS)** as dynein is not targeted and remains active.* **(a)** MSD of the MDA-MB-231 across a **+DMSO→+Blebbistatin→+Blebbistatin+Kinesore** treatment sequence reveals a substantial reduction of the cell displacement after addition of kinesin-1 activator **(+Blebb+KS)** to the Blebbistatin-pretreated cells **(+Blebb)** on both soft **(G′=8.6 kPa)** and rigid **(G′=55 kPa)** collagen grids. **(b)** While actomyosin contractility suppression **(+Blebb)** does not consistently change MDA-MB-231 cells directional ratio in comparison to the control cells **(+DMSO**, see also Figure 2b), but addition of the kinesin-1-activating Kinesore reduces directional ratio below the control baseline **(+Blebb+KS)**. **(c)** Denoised MDA-MB-231 cell migration speeds across **+DMSO→+Blebb→+Blebb+KS** sequential treatment. Suppression of the actomyosin contractility with Blebbistatin **(+Blebb)** partially slows down MDA-MB-231 cell migration velocity. Overactivation of kinesin-1 with Kinesore atop of Blebbistatin-mediated actomyosin contractility suppression **(+Blebb+KS)** further slows cell migration speeds, indicating mechanical antagonism between kinesin-1 and dynein mechanical forces during collagen-guided metastatic cell migration upon suppression of their actomyosin contractile forces. **(d)** Individual denoised migration tracks for the MDA-MB-231 cells on soft **(G′=8.6 kPa**, *top*) and rigid **(G′=55 kPa**, *bottom*) collagen grids across **+DMSO→+Blebb→+Blebb+KS** treatment sequence. **(e)** MDA-MB-231 cell migration directionalities in control **(+DMSO**, *left*), during actomyosin contractility inhibition **(+Blebb**, *center*), and during dynein co-suppression **(+Blebb+DA**, *right*), on both soft **(G′=8.6 kPa)** and rigid **(G′=55 kPa)** collagen grids. Displacement directions are compiled into circular diagrams with grid anisotropy axis oriented vertically (*arrows*).

Similarly, we examine kinesore-induced changes to contact guidance along the rhomboid collagen grid **(Figure 4d-e)**. The quantification of contact guidance shows no significant loss of cell directionality, as all MDA-MB-231 cells continue to predominantly migrate along the main grid axis on both soft (G′=8.6 kPa) and stiff (G′=55 kPa) substrates, as shown by individual cell migration tracks **(Figure 4d**, **G′=8.6 kPa** *vs.* **55 kPa, +DMSO→+Blebb→+Blebb+KS)** and total angular distribution of 1-minute-long displacement from anisotropy axis of the grid **(Figure 4e**, **G′=8.6 kPa** *vs.* **55 kPa, +DMSO→+Blebb→+Blebb+KS)**.

Therefore, kinesore-induced overactivation of kinesin-1 does not preclude dynein-driven cell adhesion, spreading, and contact guidance but attenuates the motility speed of cancer cells. Together with results from the previous experiments with direct dynein inhibition, our data show that dyneins substitute actomyosin contractility, providing locomotion and contact guidance in metastatic MDA-MB231 cells.

### Adhesion and spreading of non-metastatic cancer cells feature limited dependence on dyneins

The metastatically aggressive ^55^ and fully EMT-transitioned ^56^ triple-negative MDA-MB-231 breast cancer cells serve as one of the major models for metastasis studies ^9, 29, 30^. To further study the specific role of cytoplasmic dynein in metastasis, we choose a popular model of non-metastatic breast cancer ^57^ - a hormone receptor-positive noninvasive MCF-7 cell line, which features an incomplete epithelial-mesenchymal transition (EMT) ^58^. Presenting control MCF-7 cells **(+DMSO)** to the collagen grids results in the formation of the radial (*i.e.*, non-dendritic) protrusions that display limited adhesion restricted to the location of protrusion tips **(Figure 5a**, **+DMSO, Movie 7)**. Limited adhesion of the control MCF-7 cells prevents a complete (*i.e.*, structurally congruent) protrusions’ alignment to the collagen grids **(Figure 5b)**.

**Figure 5.**
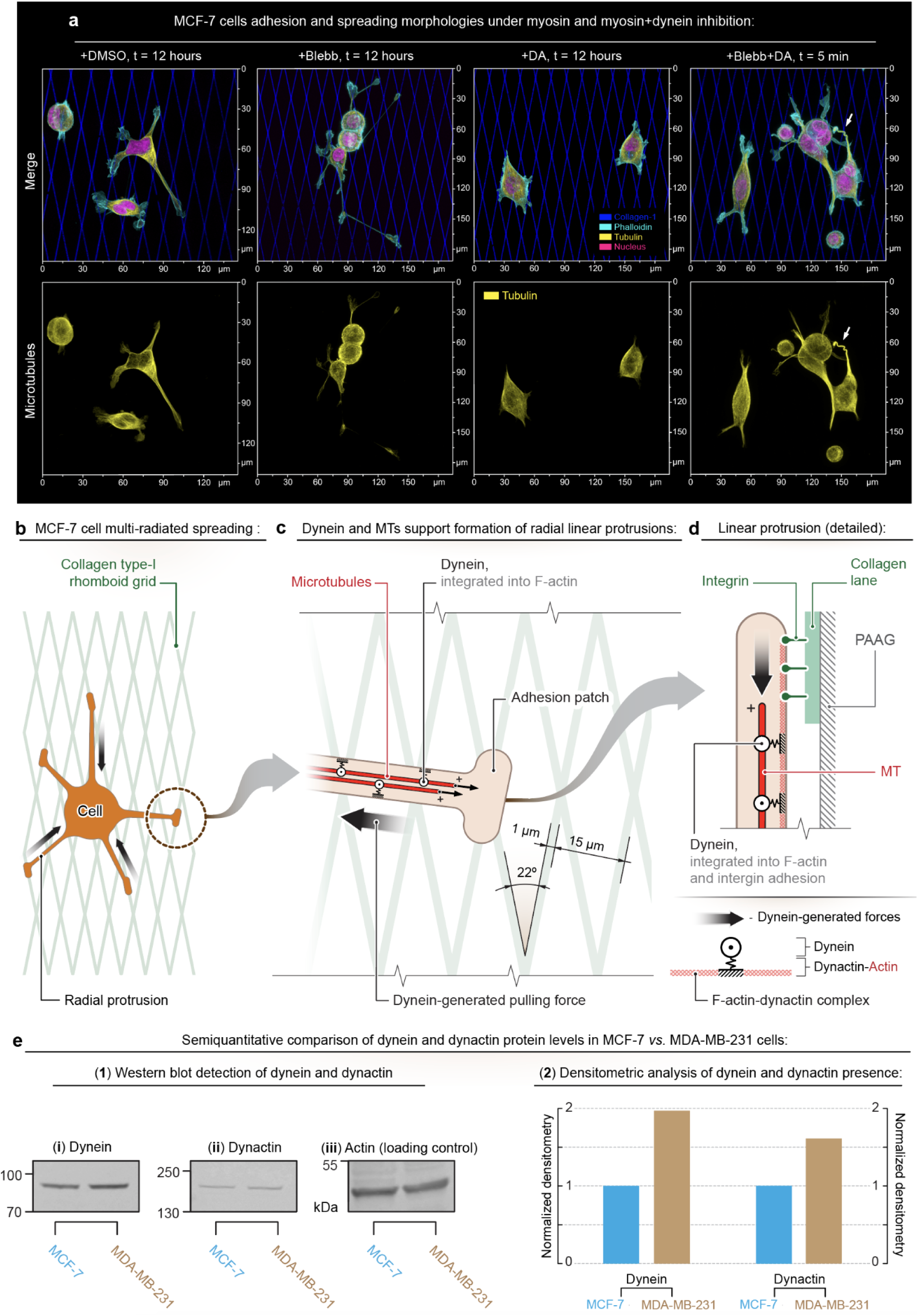
Adhesion and spreading of non-metastatic MCF-7 breast cancer cells on collagen grids display limited dependence on dynein contractility and diminished contact guidance. **(a)** Adhesion and spreading of MCF-7 cells on stiff (G′=55 kPa) collagen grids in control conditions (+DMSO), upon actomyosin contractility inhibition with Blebbistatin (+Blebb), upon dynein inhibition with Dynarrestin (+DA), and after consequential addition of Blebbistatin, and Dynarrestin (+Blebb+DA). *Note that unlike MDA-MB-231, the MCF-7 cells form radial (non-dendritic) cell protrusions with limited adhesion, restricted to their tips. Limited adhesion prevents adhesive alignment and conformity of MCF-7 protrusions to the collagen grid (+DMSO). Blebbistatin further reduces cell structural coherence by enhancing the length and number of the independent linear cell protrusions (+Blebb). Note that inhibition of dynein abrogates linear protrusions, resulting in simplified, polygonal MCF-7 cell architecture (+DA). Combined inhibition of actomyosin and dynein contractilities leads to complete cell detachment from soft (G′ = 8.6 kPa), and to partial detachment from rigid (G′ = 55 kPa) collagen-1 grids with gradual failure of the radial protrusions (arrow).* **(b)** Schematic view of the MCF-7 cell with multiple radial protrusions in control conditions (+DMSO) that do not facilitate cell alignment to the collagen grid anisotropy. **(c-d)** Suggested schematic representation of the radial protrusions-collagen grid interactions. *Note that the dynein-based structural integration of microtubules into the linear cell protrusions provides long-distance force transduction and contractility which stimulate adhesion of distal protrusion tips to the collagen grid, but not the adhesive alignment and conformity of protrusions to the collagen grid along their entire lengths.* **(e)** Semiquantitative estimation of dynein (cytosolic, clone 74.1) and dynactin-1 protein levels in MCF-7 and MDA-MB-231 cell lysates. (1) raw images of 10 μg of cell lysate, (2) graph demonstrates intensity normalized using beta actin (loading control). Densitometry of fluorescent signals indicates a 97-percent higher level of dynein and 62-percent higher level of dynactin-1 protein in MDA-MB-231 cells compared to MCF-7 line (2).

Suppression of actomyosin contractility with blebbistatin enhances the formation of the radial protrusions, reducing cells’ structural coherence, but it does not increase the protrusions’ alignment to the collagen grid **(Figure 5a**, **+Blebb)**. Alternatively, inhibition of the dyneins with dynarrestin prevents the formation of radial protrusions, inducing polygonal morphology in MCF-7 cells **(Figure 5a**, **+DA)**. These results highlight the important role of dyneins in forming radial protrusions. Abrogation of dynein’s activity in the blebbistatin-treated MCF-7 cells results in cell arrest and detachment from the soft (G′=8.6 kPa) collagen grids, while cells on the stiff collagen adhesion cues display a significant slow-down and partial detachment of the radial protrusions **(Figure 5a**, **+Blebb+DA**, *arrow*).

The limited alignment of radial protrusions to the underlying collagen grids **(Figure 5b**, **Movie 7)**, as well as limited yet substantial dependence of radial protrusions on the dynein activity **(Figure 5a**, **+DA)** highlights a lower expression of dynein and its cofactors in MCF-7, as opposed to MDA-MB-231 cell line ^59^. While limited dynein’s activity is insufficient to facilitate the alignment of cell protrusions to the collagen grids in a fully congruent manner *via* anchoring of the microtubules to the collagen, as observed in MDA-MB-231 cells **(Figure 2c)**, dynein’s activity in MCF-7 cells is still required to maintain the formation of the radial cell protrusions on collagen grid **(Figure 5c)**.

Indeed, multiple reports indicate the importance of the dynein activity ^16, 17^ and intact microtubules ^13–15, 60^ for alignment and conformational branching of cell protrusions along the adhesion cues. Similarly to the MDA-MB-231 cell model **(Figure 2c)**, we suggest that the dynein-microtubules system generates the contractility within the radial protrusions in MCF-7 cells **(Figure 5b-d)**. We argue that although mandatory, dynein contractility is only sufficient for the limited cell adhesion and morphological conformity to the adhesion cues **(Figure 5c)**, accumulating enough of the mechano-stimulatory tensile forces to activate integrins adhesion *via* the mechanosensory mechanisms at the tips of the radial protrusions ^61^. Indeed, the semiquantitative Western-blot analysis of the dynein and dynactin proteins presence in MCF-7 and MDA-MB-231 cell lines, used for the live cell motility experiments indicate substantially heightened levels of both proteins in metastatically aggressive MDA-MB-231, compared to MCF-7 cells **(Figure 5e)**.

### Limited dynein-driven spreading diminishes contact guidance of non-metastatic cells

Compared to MDA-MB-231 cells **(Figure 3a-c**, **+DMSO)**, migrating MCF-7 cells show lower cell migratory displacements (MSD) and slower locomotion in control conditions **(Figure 6a-c**, **+DMSO).** We also observe a moderate acceleration of MCF-7 cells’ speeds during inhibition of the actomyosin contractility **(Figure 5a-b**, **+DMSO)**, perhaps, due to the protrusions-dominated mode of MCF-7-collagen interactions. Thus, blebbistatin treatment is accompanied by the enhancement of the formation of the radial protrusions on both soft (G′=8.6 kPa) and rigid (G′=55 kPa) collagen grids **(Figure 6c**, **+DMSO** *vs.* **+Blebb, G′=8.6 kPa** *vs.* **55 kPa)**.

**Figure 6.**
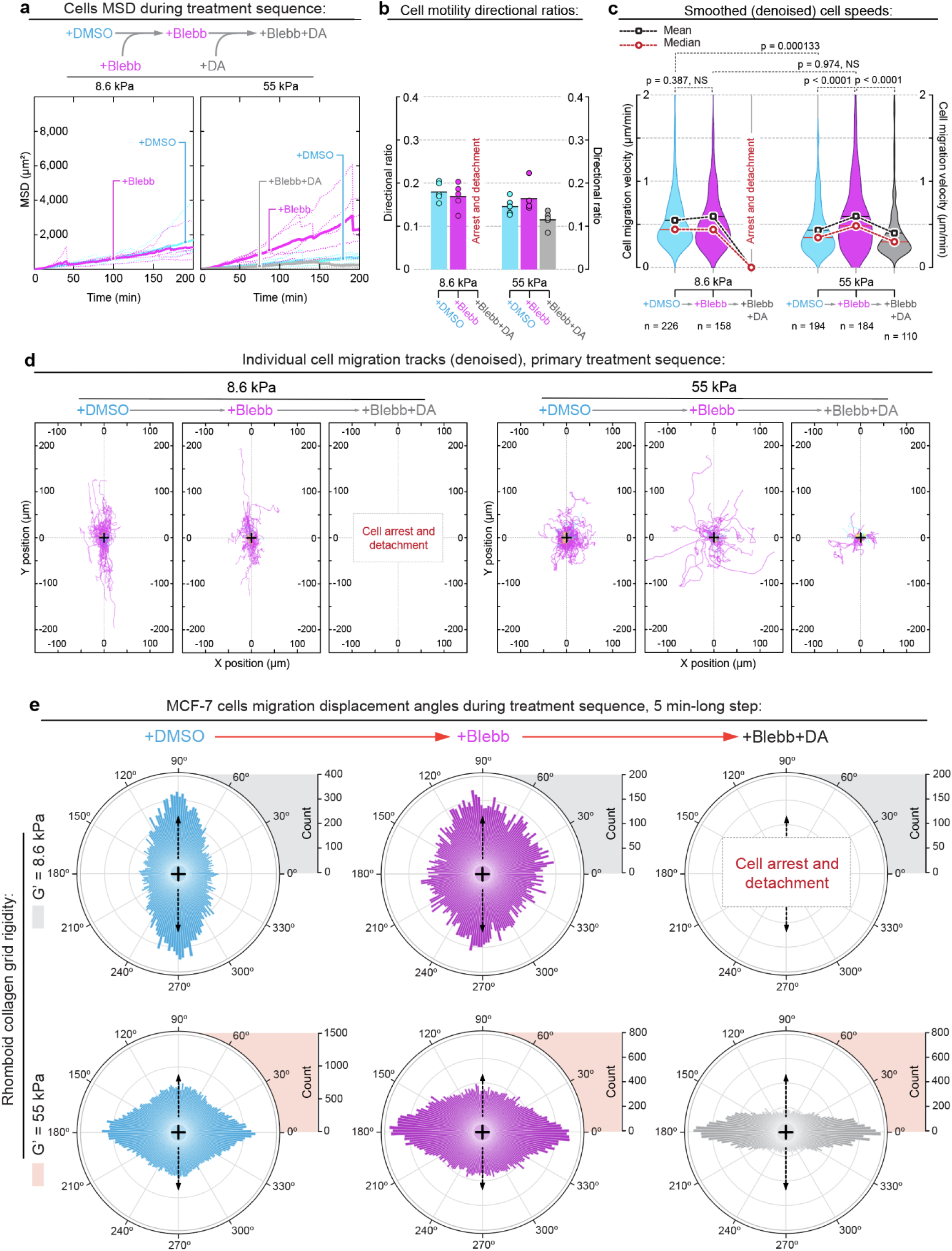
Non-metastatic MCF-7 breast cancer cells display a limited dependence on dynein activity for their adhesion and alignment to the 2D collagen contact guidance cues. **(a)** MSD of the MCF-7 cells across +DMSO→+Blebbistatin→+Blebbistatin+Dynarrestin treatment sequence displays an increase of cell displacement after addition of non-muscle myosin IIa inhibitor Blebbistatin (+Blebb) on rigid (G ’= 55 kPa) collagen grids. *Note that on soft (G′ = 8.6 kPa) collagen grids the Blebbistatin-treated MCF-7 cells arrest and detach upon dynein inhibition, while also aggregating into the clusters via E-cadherin cell-cell interactions. Note that on rigid (G′ = 55 kPa) collagen grids cells display a partial detachment and reduce their migration speeds.* **(b)** MCF-7 cells do not display a strong fluctuation of their directional ratios (*persistence*) across all conditions and rigidities. **(c)** MCF-7 migration speeds (denoised) throughout +DMSO→+Blebb→+Blebb+DA sequential treatment. Suppression of the actomyosin contractility with Blebbistatin (+Blebb) partially accelerates MCF-7 motility on both soft (G′ = 8.6 kPa) and rigid (G′ = 55 kPa) collagen grids, indicating that dynein-driven cell locomotion likely interferes with the actomyosin contractility. Further inhibition of the dyneins in addition to suppression of actomyosin contractility results in detachment of MCF-7 cells from the soft collagen grids (G′ = 8.6 kPa), while inducing a decrease of both cell speeds and directional ratio on rigid grids (G′ = 55 kPa), leading to a significant decrease of cell displacement (MSD), see panel (a). **(d)** MCF-7’s denoised migration tracks on soft (G′ = 8.6 kPa, *top*) and rigid (G ’= 55 kPa, *bottom*) collagen grids for +DMSO→+Blebb→+Blebb+DA treatment sequence. *Note a complete loss of cell migration track alignment to the stiff collagen grid (G′ = 55 kPa)*. **(e)** MCF-7 cell migration directionalities in control (+DMSO, *left*), during actomyosin contractility inhibition (+Blebb, *center*), and during dynein co-suppression (+Blebb+DA, *right*), on both soft (G′ = 8.6 kPa) and rigid (G′ = 55 kPa) collagen grids. Displacement directions are compiled into circular diagrams with grid anisotropy axis oriented vertically (*arrows*). *Note that unlike metastatic MDA-MB-231 breast cancer cells, the non-metastatic MCF-7 breast cancer cells on the rigid collagen grids (G′ = 55 kPa) display opposite response to the contact guidance cues, i.e., predominantly strides of transverse cell displacement in respect to the axis of underlying contact guidance cues. Note that most of the cell 5 min-long displacements do not lead to the effective transverse migration across the collagen grid (see panel (d), right), indicating a low efficiency of the transverse forces*.

Since MCF-7 cells feature an incomplete EMT, the underdeveloped mesenchymal organization generates insufficient actomyosin-driven ECM traction for cell migration ^62, 63^. The lack of actomyosin contractility within the cell-ECM adhesion system renders MCF-7 cells dependence on dynein+microtubules contractility within radial protrusions **(Figure 5c-d)**, confirmed by the loss of MCF-7’s radial protrusions upon inhibition of dynein **(Figure 5a**, **+DA)**. Similarly, the dependence of MCF-7 collagen interactions on dyneins is supported by the observed complete (G′=8.6 kPa) and partial detachment (G′=55 kPa) of cells upon inhibition of dyneins in blebbistatin-treated MCF-7 cells **(Figure 6c**, **+Blebb+DA)**.

As opposed to dendritic-like protrusions in MDA-MB-231 cells **(Figure 2a**, **+Blebb)**, the radial protrusions in MCF-7 cells do not sustain persistent contact guidance along the collagen grids. Compared to the anisotropy axis of the collagen grid **(Figure 6e**, **G′=8.6 kPa, +DMSO** and **+Blebb)**, individual migration tracks on soft (G′=8.6 kPa) collagen grids show relative conformity of MCF-7 cells migration along the collagen cues in control conditions **(+DMSO)** and during actomyosin inhibition **(+Blebb) (Figure 6d**, **G′=8.6 kPa, +DMSO** and **+Blebb)**, but the angular distribution of cells’ 1 minute-long displacement shows significant disarray.

Specifically, we observe a predominantly transverse orientation of MCF-7 cells’ movements with respect to the collagen grids’ anisotropy axis **(Figure 6e**, **G′=55 kPa)**. The transverse displacement without a significant change in the final cell migration track along the soft collagen grid points towards the radial protrusion-driven oscillation of MCF-7 cells across the grids. These oscillations do not result in effective cell displacement but indicate a growing role of radial protrusions in cell-collagen interactions. Moreover, these effects become substantially more evident in MCF-7 cells migrating along the rigid collagen grids (G′=8.6 *vs.* 55 kPa, **Movie 7**).

In particular, at the level of the individual cell migration track, we observe a complete loss of migration conformity in MCF-7 cells to the collagen grid in the control condition and all consequent treatments **(Figure 6d**, **G′=55 kPa, +DMSO**→**+Blebb**→**+Blebb+DA)**. We detect more frequent transversely oriented 1-minute-long displacements in relation to the collagen grid anisotropy axis **(Figure 6e**, **G′=55 kPa, +DMSO**→**+Blebb**→**+Blebb+DA)**, which become a dominant orientation for MCF-7 cells across all treatments conditions. Similarly to results on the soft collagen grids, the majority of transverse displacements are not effective, with resulting cell migration tracks displaying a random migration in relation to the underlying rigid collagen grids **(Figure 6d**, **G′=55 kPa, +DMSO→+Blebb→+Blebb+DA)**.

### Confined 3D migration of metastatic and non-metastatic cells requires combined dynein- and myosin-driven contractility

The hallmark of metastasis is the ability of cancer cells to migrate in the confinement of tissues. To test the role of actomyosin and dynein motors in the confined 3D migration, we investigate the effects of Dynarrestin **(+DA)** and Blebbistatin **(+Blebb)** on MDA-MB-231 motility in GHS. Herein, we fabricate GHS using gelatin methacryloyl (GelMA) microgel building blocks with different GelMA concentrations and varying light exposure times to provide a range of mechanical stiffnesses **(Figure S2)**.

Then, we characterize the porosity of soft (G′ ≈ 8.9±0.6 kPa) and stiff (G′ ≈ 52.2±4.8 kPa) GHS using fluorescence microscopy of scaffolds incubated in a high molecular weight fluorescent dextran solution. **(Figure 7a**, **Movies 8** and **9)**. The void fraction **(Figure 7b)**, pore identification **(Figure S3)**, and corresponding median pore diameter **(Figure 7c)** show a non-significant difference between the void fraction for soft (20±4%) and stiff (21±4%) GHS, as well as for median pore diameter: 17.8±0.8 µm and 18.3±1.0 µm for the soft and stiff GHS, respectively.

**Figure 7.**
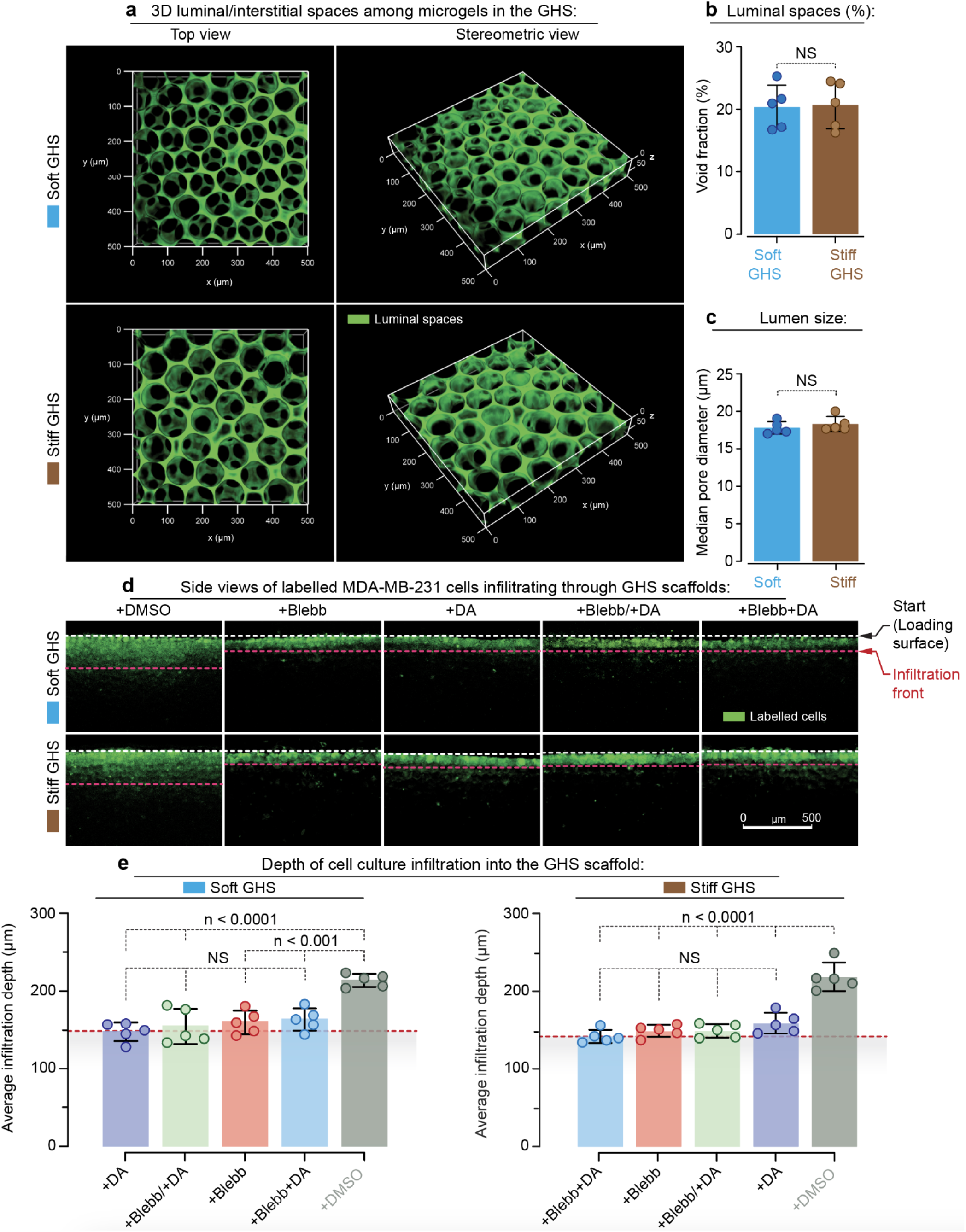
Migration of metastatic MDA-MB-231 breast cancer cells in GHS depends on simultaneous activity of dynein and myosin motors. **(a)** Three-dimensional (3D) microscopy images of GHS made up of soft or stiff GelMA microgel building blocks. The void space is occupied by fluorescently conjugated high molecular weight dextran molecules that do not penetrate the microgels. **(b)** Void fraction and **(c)** median pore diameter of soft and stiff GHS, showing no significant difference between them. **(d)** Sagittal view of sectioned scaffolds 72 h after topical cell seeding, with white and red dashed lines indicating the uppermost layer of GHS and the average cell migration depth, respectively. Cells treated with Blebbistatin **(+Blebb)**, Dynarrestin **(+DA)**, Blebb starting at *t* = 0 h, followed by DA starting at *t* = 12 h **(+Blebb/+DA)**, and both Blebb and DA starting at *t* = 0 h **(+Blebb+DA)**, showed limited motility compared with untreated cells in the control group (scale bar is 500 µm). **(e)** Average infiltrating migration length of cells treated with the inhibitors of cell contractility in GHS (*n* = 5). Dashed lines represent the average initial cell infiltration length (∼4 h after topical cell seeding) in soft (147 ± 9 µm) and stiff (138 ± 6 µm) GHS, wherein no statistically significant difference was observed compared with all the treated groups.

To study the confined 3D migration of metastatic MDA-MB231 cells, we seed fluorescently labeled cells on top of the GHS and analyze their migration for 72 h. The cross-sectional microscopy of cell-seeded scaffolds after 72 h of incubation shows no difference between control groups for soft and stiff GHS substrates **(Figure 7d** and **7e**, +DMSO). However, the confined 3D migration of MDA-MB-231 cells is lower in all treated samples **(Figure 7d** and **7e**, **+Blebb, +DA**, or **+Blebb+DA)**, as opposed to the corresponding control conditions. Moreover, the average migration length of all treated samples is limited to the extent of initial cell penetration, measured ∼4 h after seeding, where gravitational and capillary forces mainly govern cell entrainment. We observe similar results for non-metastatic MCF-7 breast cancer cells **(Figure 8a-b)** across all treated samples, which indicates that effective confined migration has more stringent requirements for simultaneous activity of myosin and dynein motors.

**Figure 8.**
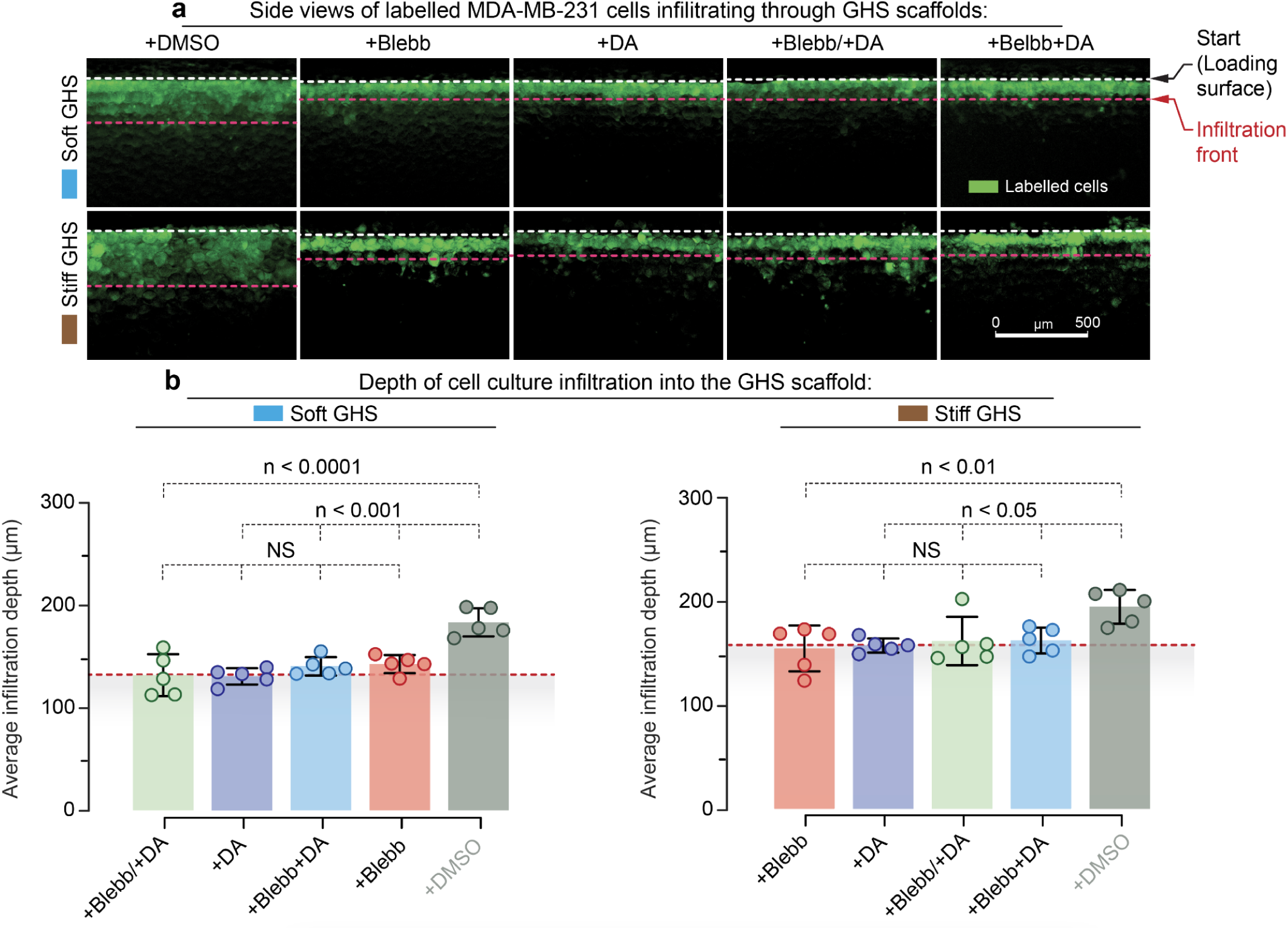
Migration of non-metastatic MCF-7 breast cancer cells in GHS depends on simultaneous activity of dynein and myosin motors. **(a)** Sectioned side view of the MCF-7 cells-infiltrated scaffolds 72 h after topical cell seeding, with white and red dashed lines indicating the uppermost layer of GHS and the average cell migration depth, respectively. Cells treated with Blebbistatin **(+Blebb)**, Dynarrestin **(+DA)**, Blebb starting at *t* = 0 h, followed by DA starting at *t* = 12 h **(+Blebb/+DA)**, and both Blebb and DA starting at *t* = 0 h **(+Blebb+DA)**, showed limited motility compared with untreated cells in the control group (scale bar is 500 µm). **(b)** Average migration length of cells treated with the inhibitors of cell contractility in GHS (*n* = 5). Dashed lines represent the average initial cell infiltration length (∼4 h after topical cell seeding) in soft (132 ± 14 µm) and stiff (160 ± 10 µm) GHS, wherein no statistically significant difference was observed compared with all the treated groups.

## DISCUSSION

The interplay between actomyosin and microtubules creates complex locomotion responses to the collagen guidance cues in metastatic MDA-MB-231 cells, as well as other mesenchymal-like cancer cells ^13, 41^. While original reports suggest that actomyosin motors are principal for cell-ECM alignment and migration within fibrillar collagen matrices ^15, 29, 64^, more recent reports indicate that actomyosin-driven cytoskeletal contractility is either insufficient or impeding the efficient motility ^16, 17, 53, 65^.

Instead, the observed morphological conformity of low contractile cells to substrate ^53^ depends both on the ‘fluid’-like polymerization dynamics of the Arp2/3-branched F-actin network and on the intact microtubule dynamics ^13, 15^. Similarly, human dermal fibroblasts switch to the low-contractility dendritic spreading in 3D collagen matrices, which relies on the presence of intact and dynamic microtubules ^66^. Moreover, the studies in fibrosarcoma cells specifically attribute the branching of low-contractile protrusions along 3D collagen matrix fibers to the dynein activity ^17^.

Indeed, dyneins mechanically slide the microtubules along the advancing axon toward the neuron’s growth cone complex ^67^. Dynein processivity within the cell cortex also mechanically repositions the entire MT apparatus and MTOC at the cell’s center ^68^. Thus, a tight mechanical and structural cooperation between the microtubules and the F-actin cytoskeleton, which is mechanically powered by dynein motors ^14, 17, 49, 67^, indicate that a similar mechanism may be utilized during metastasis.

Based on these studies, we suggest that microtubules and dyneins substitute actomyosin contractility, providing mechanical forces for cell adhesion and spreading along the collagen guidance cues, fueling the dynein-driven cancer cell metastasis. Subsequently, we show that both in metastatic mesenchymal MDA-MB-231 and non-metastatic partially mesenchymal MCF-7 cells, the migration is intertwined with the dynein-microtubules mechanics, albeit to different extents.

We suggest that in the metastatic mesenchymal breast cancer cells, dyneins and microtubules form a coherent force-generating and transmitting system that is self-sufficient to drive cell migration and contact guidance along the guidance cues of the collagen matrix. In the non-metastatic partially mesenchymal cells, dyneins facilitate cell adhesion and migration *via* the numerous yet incoherent radial protrusions. Moreover, this system provides only limited engagement of cells with the underlying collagen matrix, resulting in the lack of cell contact guidance and the low efficiency of migration in cancer cells.

In addition to the orthogonal experiment with non-metastatic breast cancer cells, we show that metastatic migration on collagen matrices depends on mutual mechanical counterbalancing of dynein and kinesin-1 microtubule-associated motors ^69^. Dynein motors display (-)-end-oriented MT processivity ^70^, while conventional kinesin-1 displays an opposite, (+)-end-oriented MT processivity ^71^. Our data indicate that regulation of dynein *vs.* kinesin-1 mechanical antagonism, *i.e.*, ‘tug-of-war’, on microtubules has a profound effect on the microtubules organization, cell contact guidance, and responsiveness to the adhesion cues through reorganization of the microtubules and their steric interactions with the surrounding microenvironments ^13, 14^.

Finally, to study metastasis in tissue-like settings, we use 3D GHS that capture multiple biomechanical aspects of native tissue. Specifically, the hydrogel scaffolds provide integrin-activation by RGD peptides, tunable local microenvironment stiffness without compromising the porosity ^72, 73^, and the confining interstitial void spaces among microgels. The interstitial void spaces enable facile nutrients, oxygen, and cellular products to exchange with the cell culture medium ^74^, resulting in a physiological-like gradient that stimulates the migration of cancer cells inside the porous scaffold ^74, 75^. As opposed to 2D contact guidance along adhesion cues, the 3D scaffolds show that migration of metastatic and non-metastatic breast cancer cells in confinement has more stringent requirements for the activity of dynein and myosin motors. Our results show that both actomyosin and dynein contractilities are required for confined migration, as upon administration of blebbistatin or dynarrestin, migration of cancer cells is restricted to initial cell infiltration length.

## MATERIALS AND METHODS

### Cell experiments

Human triple-negative breast adenocarcinoma MDA-MB-231 cells (ATCC® CRM-HTB-26™) and human breast adenocarcinoma MCF-7 cells (ATCC® HTB-26™) were cultured and proliferated in Dulbecco’s Modified Eagle Medium (DMEM), completed with 4.5 g/L D-glucose, L-glutamine,110 mg/L sodium pyruvate (Corning Cellgro®, Cat#10013CV) and 10% heat-inactivated FBS (HyClone®, Cat#SH30071.03H) at 37°C in 5% CO2. All live cell imaging and treatments were conducted in glass-bottom 35 mm cell culture dishes (MatTek Corp., Cat#P35G-1.5-14-C) using active (-)-Blebbistatin enantiomer at working concentration of 50 µM (Sigma, Cat#203391), Kinesore at 50 µM final treatment concentration (Sigma, Cat#SML2361), Dynarrestin at 50 µM cell treatment concentration (Tocris, Cat#6526), or dimethyl sulfoxide (*i.e.*, DMSO) as an empty vehicle control (Sigma, Cat#472301).

Cells were seeded at low or medium (1×10^5^-5×10^5^ cells/mL) density onto micropatterned collagen grids, mounted on the glass-bottom dishes, in the complete DMEM. For immunofluorescence imaging, cells were fixed with a fixative solution, formulated as follows: 3% Paraformaldehyde (BeanTown Chemical, Cat#50071991), 0.25% Triton X-100 (Roche, Cat#11332481001), 0.2% Glutaraldehyde (Sigma-Aldrich, Cat#340855) in 1✕PBS without Ca^2+^ and Mg^2+^ (Corning, Cat#21-040-CM) at 37°C for 15 minutes. Then samples were rinsed twice in 1✕PBS without Ca^2+^ and Mg^2+^ for 10 minutes each cycle, then quenched in cold (*i.e.*, t = 4°C) cytoskeletal buffer (*i.e.*, 10 mM MES (Sigma-Aldrich, Cat#M3671-50G), 150 mM NaCl (Flinn Scientific, Cat#S0062), 5mM EGTA (Sigma-Aldrich, Cat#4100-50GM), 5 mM MgCl_2_ (Fisher Scientific, Cat#M33-500), 5 mM Glucose (Acros, Cat#41095-0010), pH 6.1) with freshly added sodium borohydride (1 mg/mL; TCI, Cat#S0480) for 15 minutes on ice. The quenched samples were then rinsed three times in 1✕PBS, 5 minute each cycle, then blocked with 1% BSA (Fisher Bioreagents, Cat#BP9704-100) and immunostained ^76^. For immuno-fluorescent microtubules staining, we utilized primary antibodies against β-tubulin (Sigma, Cat#T7816) diluted in 1% BSA PBS, incubated with the samples for 1 hour at ambient temperature of 20°C. Consequently, primary antibodies-labeled structures were stained with Alexa-Fluor-conjugated secondary antibodies (Thermo Fisher) at the final concentration of 5 µg/mL, 1 hour in 1% BSA PBS at ambient temperature. F-actin was selectively labeled with fluorescent phalloidin (Phalloidin-ATTO 647N conjugate, Millipore-Sigma, Cat#65906; 10 U/mL). Cell chromatin was labeled with 1:1000 Hoechst solution (Tocris, Cat#5117). Samples were mounted using 90% Glycerol (Sigma, Cat#G5516) in 1✕ PBS.

### High precision micropatterning

A step-by-step methodological instruction and a detailed protocol for high-fidelity, sub-micron precision polyacrylamide gels micropatterning with various ligands, to include collagen type-1, is described elsewhere ^77^. Briefly, in order to prevent van-der-Waals and common deformation effects that routinely cause the collapse of soft lithographic stamp (*i.e.*, soft regular PDMS (rPDMS)) onto the printed glass surfaces, we replaced rPDMS with a composite - *i.e.*, soft cushioning rPDMS layer, veneered with a ∼0.5 mm hard non-collapsing PDMS (hPDMS, see “hPDMS formulation” section) ^13, 78^. We cast micro-printing surfaces with the commercially premade silicon crystal molding matrix (Minnesota Nano Center, University of Minnesota) by coating the passivated molds with ∼0.5 mm layer of hPDMS, followed by 30 minutes-long curing at 70°C. Next, an 5-6 mm-thick layer of degassed rPDMS liquid premix is poured atop of the solidified hPDMS layer (rPDMS; 1:5 curing agent/base ratio, Sylgard-184, Dow Corning), then baked at 70°C for ~1 hour. The resulting cured composites were then released from the molding surfaces, and carefully cut into square blocks of 1✕1 cm in size.

Collagen type-1 protein is not directly printable *via* the means of soft micro-contact lithography due to its gelation. Thus, to prepare the collagen type-1 micro-patterns, we microprinted anti-collagen type-1 polyclonal rabbit Ab (AbCam, Cambridge, UK, Cat#ab34710; RRID:AB_731684), pre-conjugated with a biotin tag, ((+)-biotin *N*-hydroxysuccinimide ester, Sigma-Aldrich, Cat#H1759; commercial protocol) and a fluorescent tag (Alexa Fluor® succinimidyl esters, Invitrogen™, Molecular Probes®, Cat#A20000, Cat#A20003; as per supplied protocol) on a clean (*i.e.*, dust-free) intermediate coverglass (FisherFinest™ Premium Cover Glass; #1, Cat#12-548-5 P). Soft lithographic micro-printing was executed by incubating the composite micro-stamp’s surfaces with the 0.2 mg/mL biotin and fluorophore pre-conjugated α-collagen-1 antibody, reconstituted in 1✕PBS (40 min, 37°C) in a humid chamber. The coated stamps were rinsed in deionized water, then blow-dried with air. The dried stamps were then gently applied to the intermediate glass surfaces with their antibody-coated printing surfaces, then 100 g weight was placed atop the stamp to ensure proper contact between the stamping surface and the intermediate glass for 1 minute and then removed. The quality of the resulting micropatterns was examined with the epifluorescent microscope (ThermoFisher, EVOS M5000, Cat#AMF5000).

35 mm cell culture plastic dishes with 20 mm circular cutouts, sealed with the glass bottom (MatTek Corp., Ashland, MA, Cat#P35G-1.0-20-C) were chemically activated to covalently cross-link with polyacrylamide (PAA) gels, *i.e.*, coated with 3-(trimethoxysilyl)propyl methacrylate (Sigma-Aldrich, Cat#6514, as per commercial protocol). Small volumes (*e.g.*, ∼5 μL) of PAA premix (see the “PAA elastic gel premix” section) of the projected rigidity G′ = 8.6 or 55 kPa ^79^, complemented with 5% streptavidin-acrylamide (ThermoFisher, Cat#S21379), were ‘sandwiched’ between the activated dish’s glass bottom and the micropatterned intermediate glass upon adding a curing catalyst (aminopropyltriethoxysilane (APS)). After PAA curing is complete (∼5 minutes), the resulting PAA ‘sandwiches’ were incubated in hypotonic (*i.e.*, deionized) water for 1 hour at room temperature (20°C) to ensure PAA gel osmotic swelling and its gentle deformation for easier intermediate glass release from the PAA layer. The resulting PAA gels, attached to the glass bottoms and with the cross-linked antibody micropatterns, were quality-controlled and selected by examining the micropatterns’ quality with an epifluorescent microscope. The selected micropatterns with α-collagen-1 Ab were then incubated with 1 mg/mL rat monomeric collagen-1 (Corning, NY, Cat#354249) in cold PBS (4 °C, 12 hours), then rinsed three times with cold PBS, and utilized for experiments.

### hPDMS formulation

hPDMS premix composition is as follows: 3.4 g of VDT-731 (Gelest, Inc., Cat#VDT-731), 18 μL of Pt catalyst (Platinum(0)-2,4,6,8-tetramethyl-2,4,6,8-tetravinylcyclotetrasiloxane complex solution) (Sigma-Aldrich, Cat#479543), 5 µL of cross-linking modulator 2,4,6,8-Tetramethyl-2,4,6,8-tetravinylcyclotetrasiloxane (Sigma-Aldrich, Cat#396281). The resulting mixture was mixed in a 50 mL conical bottom centrifuge tube on the vortex mixer for at least 30 sec, then 1 g of HMS-301 (Gelest, Inc., Cat#HMS-301) was added immediately before use and mixed for 30 sec on a vortex mixer ^78, 80^.

### PAA elastic gel premix

To prepare the customized PAA premixes with projected gel rigidity, we utilized 40% acrylamide (40% AA) as a base (BioRad, Cat#161–0140), and 2% bis-AA (BioRad, Cat#161–0142) as a cross-linker ^81, 82^. The PAA premixed was also supplemented with streptavidin-acrylamide (Thermo Fisher, Cat#S21379) to the final concentration of 0.133 mg/mL to facilitate PAA gels cross-linking with biotin-conjugated anti-collagen antibodies (*i.e.*, micropatterns). For the preparation of 5 µL of G′ = 8.5 or 55 kPa PAA gel premixes respectively, the components were mixed as follows: 40% AA - 2.34 or 2.25 µL; 2% bis-AA - 1.88 or 2.25 µL; 2 mg/mL streptavidin-acrylamide - 0.33 µL for both rigidities; 10✕PBS - 0.5 µL for both rigidities; deionized milli-Q water - 1.117 µL for both rigidities; TEMED - 0.01 µL for both rigidities. The premix solutions were ultrasonicated to remove gas, then stored at 4°C before use. To initiate the polymerization of PAA premixes 0.1 µL of 10% APS was added to 5 µL of PAA premix immediately prior to the PAA gel casting procedure.

### Western blot

For cell extract preparation, confluent MCF-7 and MDA-MB 231 cells were washed twice with PBS. A mixture of RIPA Buffer-2 (2X, ThermoScientific,Cat#J60629EQE) and cOmplete™, Mini, EDTA-free Protease Inhibitor Cocktail (Roche Diagnostics, Cat#11836170001) was added to uniformly cover the dish surface. The dishes were then incubated on ice for 10 minutes. After, the cell extract was scraped off the dish and collected in a microcentrifuge tube, sonicated for 30 seconds at 50% pulse, and centrifuged at 21000 x g for 15 minutes. Following centrifugation, the supernatant was collected and stored at -20°C. The Pierce™ BCA Protein Assay Kit (Cat#23227) was used to determine the protein concentrations of the cell extracts. For use in Western Blot, a solution of 1X NuPAGE™ LDS Sample Buffer (4X, Invitrogen, cat no. NP0008), 1X NuPAGE™ Sample Reducing Agent (10X, Invitrogen, Cat#NP0009), and 1 µg/µL cell extract was prepared. This solution was warmed at 100°C for 10 minutes then stored at -20°C.

After removing the cell extract solution from the freezer, the samples were heated at 70°C for ten minutes. The 10 μg of samples and the PageRuler Plus Prestained Protein Ladder (Protein Biology, Cat#26619) were then loaded onto a NuPAGE™ 4-12% Bis-Tris Gel with 12 wells (Invitrogen). The gel was run with 1X NuPAGE™ MOPS SDS Running Buffer (20X, Invitrogen, Cat#NP0001-02) and ran at 150 V for 75 minutes. The gel was then transferred to low-fluorescence PVDF transfer membrane (ThermoFisher), which was placed into iBlot® PVDF Mini Stack, replacing the PVDF membrane that was pre-packaged into stack. Next, the Mini Stack was moved into the iBlot®2 instrument, where the protein transfer occurred at 20 V for seven minutes. Following the transfer, the membrane was blocked in Intercept® TBS Blocking Buffer (LI-COR) for four hours, shaking at room temperature.

After blocking, the membrane was cut into three sections, one for each protein (Cell Signaling). Each membrane piece was then incubated overnight at 4 °C, shaking, with the target protein’s respective antibody. The following primary antibodies were used: mouse anti-dynactin 1 (1:250, BioLegend, Cat#867602), mouse anti-dynein (1:1000, Millipore, Cat#MAB1618), and mouse anti-beta actin (1:1000, Abcam, Cat#ab8226). The antibodies were diluted with Intercept® Antibody Diluent T20 TBS (LI-COR, Cat.#927-65001). After overnight incubation, the membranes were washed with 0.05% TBS-T three times, five minutes each, shaking at room temperature.

Next, the membrane pieces were incubated for one hour at room temperature, shaking, with the respective secondary antibody. For the mouse-isotype primary antibodies, IRDye® 680RD Goat anti-Mouse Secondary Antibody (1:20000, LI-COR, Cat#926-68070) was used. Antibody was diluted with the Intercept® Antibody Diluent T20 TBS. Following the incubation, the membranes were again washed with 0.05% TBS-T three times, five minutes each, shaking at room temperature. The membranes were then imaged on a LI-COR Odyssey CLx and analyzed via ImageStudio (version 5.2).

## 2D cell migration assay

The confocal imaging was performed on the Leica TCS SP8 laser scanning confocal microscope with LIAchroic Lightning system and LAS X Lightning Expert super-resolution capacity, 405, 488, 552 and 638 nm excitation diode lasers, with 40✕/1.3 oil immersion objective (Leica, Germany). The scanning settings were optimized with Nyquist LAS X function with HyD2.0-SMD excitation sensors, at the regular pinhole size of 0.85 AU, and the scanning frequency at 100 Hz. Each frequency channel was scanned in sequence in order to avoid signal bleeding between the channels. Morphometric analysis was performed automatically and/or manually utilizing LAS X (Leica, Germany) and ImageJ/FIJI. Figures were composed using Adobe Illustrator CC 2017 (Adobe).

### GelMA synthesis

GelMA synthesis was conducted based on our previously established protocol ^72, 73^. Briefly, 2.5 mL of methacrylic anhydride (Sigma, MO, USA) was added to DPBS (Gibco, MA, USA) containing 10% w/v of porcine skin gelatin powder (∼300 g Bloom, Type A, Sigma, MO, USA) dropwise and allowed to react for 2 h at 50 °C while stirring. Then, DPBS was added to the solution (2:1 volume ratio) to stop the reaction. The solution was dialyzed for 10 days against ultrapure Milli-Q^®^ water (Millipore Corporation, MA, USA) at 40°C using standard grade regenerated cellulose dialysis membranes (Spectra/Por^®^ 4 with Mw cutoff = 12-14 kDa, Spectrum Laboratories, NJ, USA). The final solution was then filtered using vacuum filtration (pore size = 0.2 µm, VWR, PA, USA). The GelMA solution was then transferred to 50 mL centrifuge tubes (Celltreat, MA, USA) and subsequently frozen at -80°C while being placed in a sideways position. After 48 h, the frozen GelMA was lyophilized for 3 days to yield solid GelMA.

### Bulk hydrogel scaffold fabrication

Bulk hydrogel scaffolds were fabricated using different GelMA concentrations and varying light exposure times. Soft or stiff scaffolds were prepared by dissolving GelMA in a photoinitiator solution (0.1% w/v of lithium phenyl-2,4,6-trimethylbenzoylphosphinate (LAP, Sigma, MO, USA) in DPBS) at 37 °C at the concentrations of 50 mg mL^-1^ or 150 mg mL^-1^, respectively. The dissolved GelMA was then poured into a cylindrical mold (diameter = 8 mm and height = 1 mm). The molded GelMA solution was then placed in a humidity chamber made from a petri dish with wet Kimwipes and stored at 4 °C overnight to form a physically crosslinked gel while protected from light. Finally, physically crosslinked molded GelMA hydrogels were exposed to light (wavelength = 395-405 nm and intensity = 15 mW cm^-2^) for 1 min (5% w/v GelMA solution) or 2 min (15% w/v GelMA solution) to yield the soft or stiff scaffolds.

### GelMA microgel fabrication

A step emulsification microfluidic device was fabricated for high throughput microgel production as described previously ^72, 73^. For soft or stiff GelMA microgels, 50 mg mL^-1^ or 150 mg mL^-1^ of GelMA solution was prepared in 01% w/v photoinitiator solutions, respectively. Novec™ 7500 engineered fluid (3M, MN, USA) supplemented with Pico-surf™ (2% v/v) (Sphere Fluidics, Cambridge, UK) was prepared for the continuous phase. The GelMA and Pico-surf™ solutions were then injected into the microfluidic device using syringe pumps (PHD 2000, Harvard Apparatus, MA, USA). The temperature was maintained at 35-40 °C using a space heater. Once collected in microcentrifuge tubes, microgel suspensions were stored at 4 °C to physically crosslink.

### GHS fabrication

While protected from light, oil and surfactant were removed from the microgel suspension using 1H,1H,2H,2H-perfluoro-1-octanol (20% v/v, Alfa Aesar, MA, USA) in Novec™ 7500 engineered fluid. To this end, the solution was added to the suspension (1:1 volume ratio) and vortexed for 10-20 s, followed by centrifugation at 325 × g for 15 s. Then, 400 µL of photoinitiator solution (LAP in DPBS, 0.1% w/v) was added to the suspension, vortexed for 10 s, and centrifuged at 325 × g. This step was followed by the addition of 400 µL of photoinitiator solution, vortexing for 10 s, and centrifugation at 1310 × g. The supernatant was decanted, and microgel suspension was pipetted into a mold (diameter = 10 mm and height = 3 mm), using a positive displacement pipette (Microman E M100E, Gilson, OH, USA). The molded microgels were then placed in a chamber and exposed to light (395-405 nm and 15 mW cm^-2^) for 1 min or 2 min to form GHS made up of soft or stiff microgels (termed as soft GHS or stiff GHS), respectively.

### Rheological characterizations

The rheological properties (storage and loss moduli) of bulk hydrogel scaffolds, representing the individual microgels, were characterized by placing the scaffolds (diameter = 8 mm and height = 1 mm) between two sandblasted parallel plates (upper plate diameter = 8 mm, bottom plate diameter = 25 mm) in an AR-G2 rheometer (TA instrument, DE, USA). Once sandwiched between the plates, the amplitude sweep test was conducted on samples at a constant frequency of 1 rad s^-1^ to find the linear viscoelastic region (LVR). Then, a frequency sweep test was performed on samples at a constant oscillatory strain of 0.1% (within the LVR) from 0.1 to 100 rad s^-1^.

### Pore characterization of GHS

GHS were incubated in a fluorescein isothiocyanate–dextran solution (Sigma, MO, USA, with an average molecular weight of 2 MDa) in Ultra-Pure Milli-Q^®^ water (15 µM). The scaffolds were then imaged using a Leica DMi8 Thunder™ microscope (Germany). The Leica application suite X (LAS X, version 3.7.4.23463) software was used to construct 3D images from high-magnification (250×) *z*-stacked images (141 z-slices, increment size = 0.502 µm). The void fraction was then measured as the fraction of occupied void space over the total volume, using LAS X software. Also, for pore detection and reporting equivalent median pore diameter, a MATLAB code was developed and used, wherein 2D fluorescence microscopy images were thresholded, and the pores were detected and masked. The areas of detected pores were obtained and converted to equivalent-area circles, from which the diameters were calculated. The median of the equivalent diameter distribution was reported.

### 3D cell migration assay

Cell migration was analyzed using the MDA-MB-231 or MCF-7 cell lines labeled with CellTracker™ Green 5-chloromethylfluorescein diacetate (CMFDA) Dye (Invitrogen, MA, USA). Briefly, 50 µg of the dye was reconstituted in 20 µL of DMSO (Sigma. MO, USA). Then, the dye solution was diluted in DMEM (Gibco, MA, USA) (1:1000 volume ratio), and 10 mL of dye-containing culture media was added to the cell culture flasks (VWR, PA, USA). Cells were incubated under a 5% v/v CO_2_ atmosphere at 37 °C for 30 min. Then, culture medium was replaced with DMEM, supplemented with 10% v/v FBS (Cytiva, MA, USA) and 1% v/v antibiotic (10000 U mL^-1^ penicillin G, 10000 µg mL^-1^ streptomycin, 25 µg mL^-1^ amphotericin B, Cytiva, MA, USA). After 3 h of incubation in antibiotic-containing DPBS (1% v/v) at room temperature, scaffolds were transferred to a non-treated 24-well cell culture plate (VWR, PA, USA). Then, 20 µL of cell suspension (5×10^6^ cells per mL in complete cell culture media) was added on top of the scaffolds. After topical seeding, scaffolds were incubated at room temperature for ∼30 min, and then complete cell culture media with or without cell contractility inhibitors (final working concentration of 50 µM) were added to each well, and the culture dish was placed in an incubator (5% v/v CO_2_ atmosphere at 37 °C). After 72 h, samples were sectioned through their sagittal plane, using a razor blade (VWR, PA, USA) and imaged using the Leica DMi8 Thunder™ microscope at an excitation wavelength and emission wavelength of 470 nm (blue) and 550 nm (green), respectively. Then, cell migration length was measured throughout the scaffold at different points using the ImageJ software (FIJI, version 1.53t, NIH, MD, USA) ^83^, and the average migration length was reported.

### Quantification of cell migration on anisotropically aligned collagen patterns

The quantification analyses for Figures 3, 4, 6, and S1 were performed with an automated pipeline based on in-house MATLAB scripts. The pipeline includes the following steps.

To identify individual cells in time-lapsed-based phase contrast images, we: (1) evened the background by subtracting images smoothed by Gaussian kernel with a specified standard deviation (20 pixels for our 2048x2048 data) from the original images; (2) identified local variations (cells, cell edges, and protrusions) with the standard deviation (STD) filter using 5-by-5 neighborhood; (3) segmented the resulting STD map; (4) corrected for over-segmentation by the erosion filter using a circular structural element of radius 15 pixels; (5) filtered out very small and very large objects (< 10 and >7000 pixels, respectively); (6) recorded the coordinates of the centroids of the remaining objects (cells). To choose the proper threshold values for step (3), we run an optimization routine that finds a threshold value maximizing the number of identified cells in each image at the end of the processing.

Next, we used a pipeline for extracting cell tracks. This pipeline consists of (1) matching cells between consecutive time frames; (2) correcting for flickering events; and (3) correcting for track cloning. In the first step, we match each cell at time *t* with the nearest cell at time *t*+1 within a circle of radius *R*_max_ (20 pixels in for our data). Because cells in our data may have very complex geometries with cell bodies being on the edge of the algorithm’s detection margins, it is possible that tracks become interrupted by cell ‘disappearance’ for one time point (a.k.a. flickering). In step (2), we use an algorithm that detects such events based on the proximity of tracks ending at time *t* and tracks starting at time *t*+2 and connects the interrupted tracks together. For the same shape complexity reason, it is possible that occasionally one cell can be detected as two closely located ones. In such cases, we can obtain two trajectories representing one cell and closely following each other. In step (3), we use an algorithm that detects such cloned tracks and eliminates them from the record.

In our data, the detected tracks accurately match the observed complex movement of the cells. However, frame-to-frame variations in cell shape and slight jiggling of the microscope stage contribute to a pixel-size noise along the cell tracks. To minimize such instantaneous distortions, we smoothed all tracks by applying running average to *x* and *y* coordinates of the track points (15 min window in 1 min steps) before calculating angular directions and velocity values along the tracks.

### Statistical analysis

We utilized the pairwise one-sided *t*-tests between a control group and corresponding perturbation conditions. Computation of the statistical data was performed with KaleidaGraph 4.5.4 (Synergy Software) and Prism 7b (GraphPad Software, Inc). The numeric p-values are denoted on the corresponding plots. If p-value is below 0.0001, *i.e.*, cut-off lower limit for Kaleidagraph and Prism 7b, a ‘p<0.0001’ is denoted. Sample size n (*i.e.*, number of analyzed cells) for each measured parameter is denoted on the plots. Each measured parameter is a result of three replicates (independent experiments), unless specified otherwise. Box and whiskers plots reflect the first quartile, median, third quartile, and 95% percent confidence interval. Data shown as violin diagrams are distributions. Mean and median values are outlined separately.

## CONFLICTS OF INTEREST

There are no conflicts of interest to declare.

## DATA AVAILABILITY STATEMENT

The authors declare that the data supporting the findings of this study are available within the paper, the supplementary information, and any data can be made further available upon reasonable request. The raw data and the code used in analysis, cell identification and tracking pipeline will be available through GitHub depository (https://github.com/tsygankov-lab).

## ACKNOWLEDGEMENTS

E.D.T., Y.T. and this work were supported by the Department of Pharmacology, Penn State College of Medicine *via* the startup funds. This research received funding from the Meghan Rose Bradley Foundation (A.S.). A.S. would like to acknowledge Penn State startup fund. This work was also supported by grants from the National Science Foundation (CMMI 1942561) and the National Institutes of Health (R01GM136892) to D.T.; A.N., O.P., X.M., A.S.Z would like to acknowledge Intramural FDA funding.

## Movie legends

**Movie 1.** MDA-MB-231 cell migration and morphology in control conditions **(+DMSO**, G′=55 kPa). Collagen type-1 rhomboid grid is partially depicted in a single quadrant of the field of view. Scale bar and timestamp are denoted on the video.

**Movie 2.** MDA-MB-231 cell migration and morphology during low actomyosin contractility **(+Blebb**, G′=55 kPa). Scale bar and timestamp are denoted on the video.

**Movie 3.** Failure of the blebbistatin-induced dendritic-like protrusions and MDA-MB-231 cells detachment upon inhibition of the dynein with dynarrestin **(+Blebb→+Blebb+DA**, G′=55 kPa). Scale bar and timestamp are denoted on the video.

**Movie 4.** Compiled video-sequences for migration of MDA-MB-231 cells along soft (G′=8.6 kPa, *left*) and rigid (G′=55 kPa, *right*) collagen type-1 rhomboid grids during primary treatment sequence **(+DMSO→+Blebb→+Blebb+DA)**. The control condition **(+DMSO***, top*) and low actomyosin contractility **(+Blebb***, bottom*) cell migration state are shown. Cell migration tracks are highlighted as a computed cell centroid displacement (*yellow tracks*). See cell detachment in **(+Blebb+DA on** G′=55 kPa) on Movie 3. Frame frequency (*i.e.*, frames per second, fps) is 32 fps, each frame corresponds to 1 minute of real time microscopy. Scale bar is 20 µm on the panels with denoted tracks. Scale bar on the zoom-in panels with stationary cell centroids is 50 µm.

**Movie 5.** Video-sequences for MDA-MB-231 cells migrating along the soft (G′=8.6 kPa, *left*) and rigid (G′=55 kPa, *right*) collagen grids, reversed treatment sequence **(+DMSO→+DA→+DA+Blebb)**. Control cells **(+DMSO***, top*) and inhibited dynein **(+DA***, bottom*) cell migration states are shown. Cell migration tracks are highlighted as a computed cell centroid displacement (*yellow tracks*). Frame frequency (*i.e.*, frames per second, fps) is 32 fps, each frame corresponds to 1 minute of real time microscopy. Scale bar is 20 µm on the panels with denoted tracks. Scale bar on the zoom-in panels with stationary cell centroids is 50 µm.

**Movie 6.** Compiled migration modes sequences for migrating MDA-MB-231 cells along the soft (G′=8.6 kPa, *left*) and rigid (G′=55 kPa, *right*) collagen grids during **+Blebb→Blebb+KS** treatment sequence. Low actomyosin contractility conditions alone **(+Blebb***, top*), and its combination with enhanced kinesin-1 activity **(+Blebb+KS***, bottom*) cell migration modes are shown. Cell migration tracks are highlighted as a computed cell centroid displacement (*yellow tracks*). Frame frequency (*i.e.*, frames per second, fps) is 32 fps, each frame corresponds to 1 minute of real time microscopy. Scale bar is 20 µm on the panels with denoted tracks. Scale bar on the zoom-in panels with stationary cell centroids is 50 µm.

**Movie 7.** Live video-sequences for MCF-7 cell migration along the soft (G′=8.6 kPa, *left*) and rigid (G′=55 kPa, *right*) collagen type-1 grids. The control conditions **(+DMSO***, top*) and low actomyosin contractility **(+Blebb***, bottom*) cell migration states are shown. Cell migration tracks are highlighted as a computed cell centroid displacement (*yellow tracks*). See cell detachment in **(+Blebb+DA on** G′=55 kPa) on Movie 3. Frame frequency (*i.e.*, frames per second, fps) is 32 fps, each frame corresponds to 1 minute of real time microscopy. Scale bar is 20 µm on the panels with denoted tracks. Scale bar on the zoom-in panels with stationary cell centroids is 50 µm.

**Movie 8.** 3D microscopy reconstruction of soft (G′ ≈ 8.9 ± 0.6 kPa) GHS scaffold (interstitial space is green).

**Movie 9.** 3D microscopy reconstruction of stiff (G′ ≈ 52.2 ± 4.8 kPa) GHS scaffold (interstitial space is green).

---

**Supplementary Figure S1.**
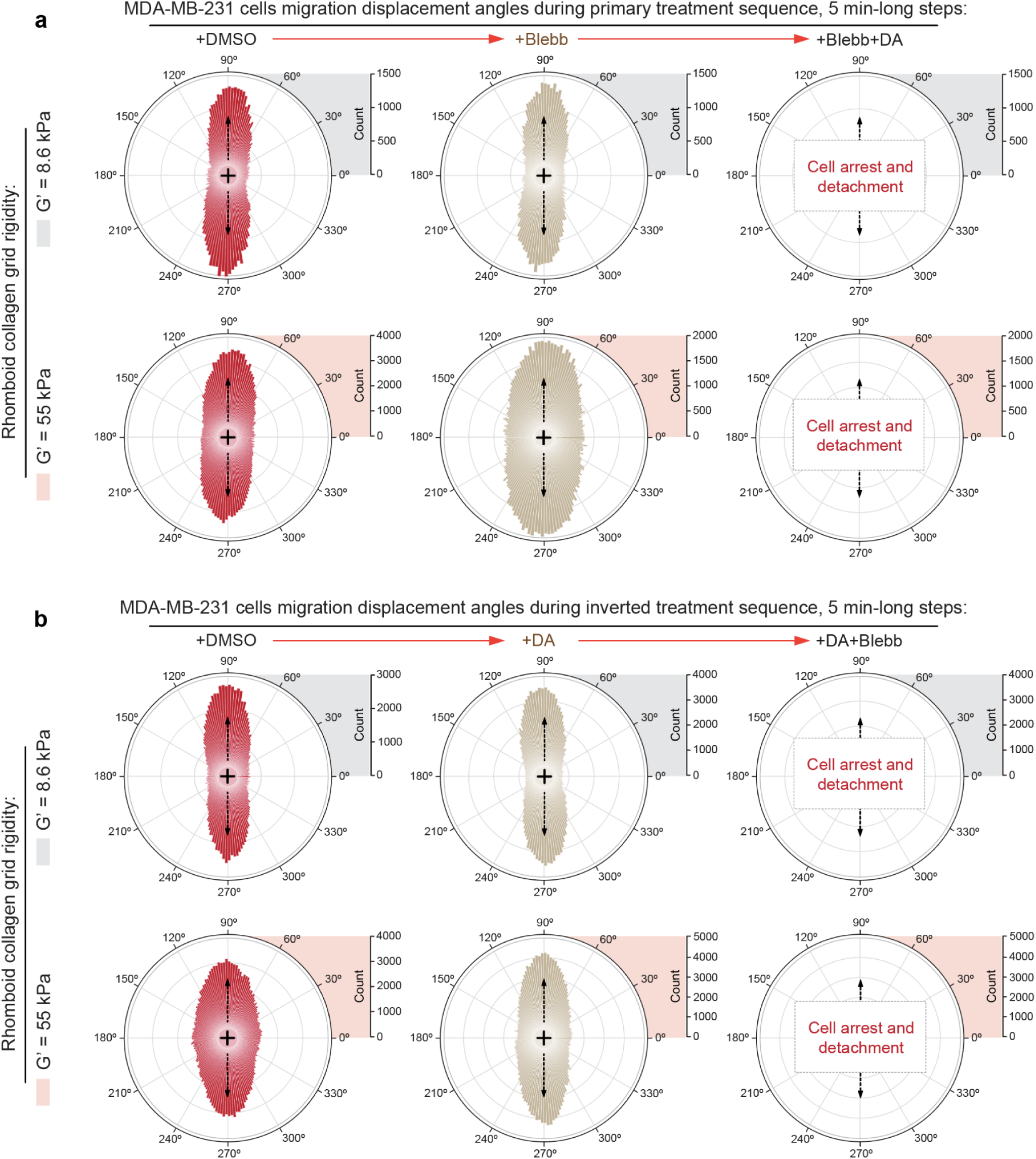
Distribution of angular deflections for cell migratory displacements from anisotropy axis of the collagen rhomboid grid for metastatic MDA-MB-231 and non-metastatic MCF-7 breast cancer cell lines in various conditions. Cumulative directions distribution of individual cell displacements are calculated for 5 minute-long timeframe steps. Displacement directions are compiled into circular diagrams with grid anisotropy axis oriented vertically (*arrows*). **(a)** MDA-MB-231 cell migration directionalities in control conditions **(+DMSO**, *left column*), followed by the actomyosin contractility inhibition **(+Blebb**, *central column*), concluded by dynein co-suppression *via* adding Dynarrestin to the Blebbistatin-treated cells **(+Blebb+DA**, *right column*), on soft **(G′ = 8.6 kPa**, *top row*) and rigid **(G′ = 55 kPa**, *bottom row*) rhomboid collagen grids. (b) Directionalities of MDA-MB-231 cell displacements during inverted treatment sequence: in control conditions **(+DMSO**, *left*), followed by dynein activity inhibition **(+DA**, *center*), with subsequent actomyosin contractility co-inhibition along with continued dynein suppression **(+DA+Blebb**, *right*).

**Supplementary Figure S2.**
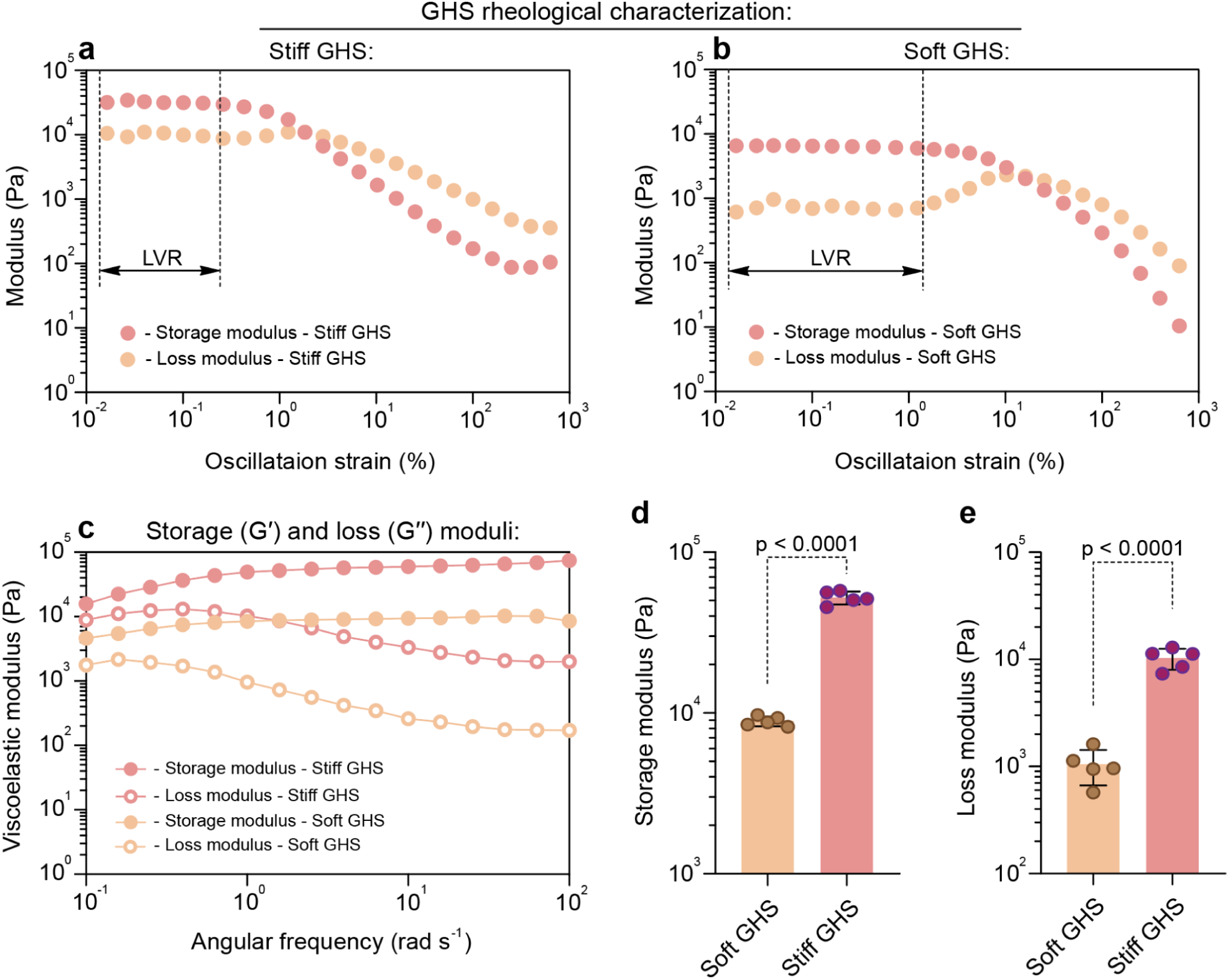
Rheological characterization of GHS. Oscillation strain sweep of **(a)** Stiff or **(b)** Soft GHS, measured at a constant frequency of 1 rad s^-1^ to identify the linear viscoelastic region (LVR). **(c)** Storage (*G*′) and loss (*G′′*) moduli of soft and stiff GHS versus angular frequency, measured at an oscillatory strain of 0.1%. **(d)** Average storage modulus of soft and stiff GHS acquired at a strain of 0.1% and angular frequency of 1 rad s^-1^. **(e)** Average loss modulus of soft and stiff scaffolds measured at a constant strain of 0.1% and angular frequency of 1 rad s^-1^.

**Supplementary Figure S3.**
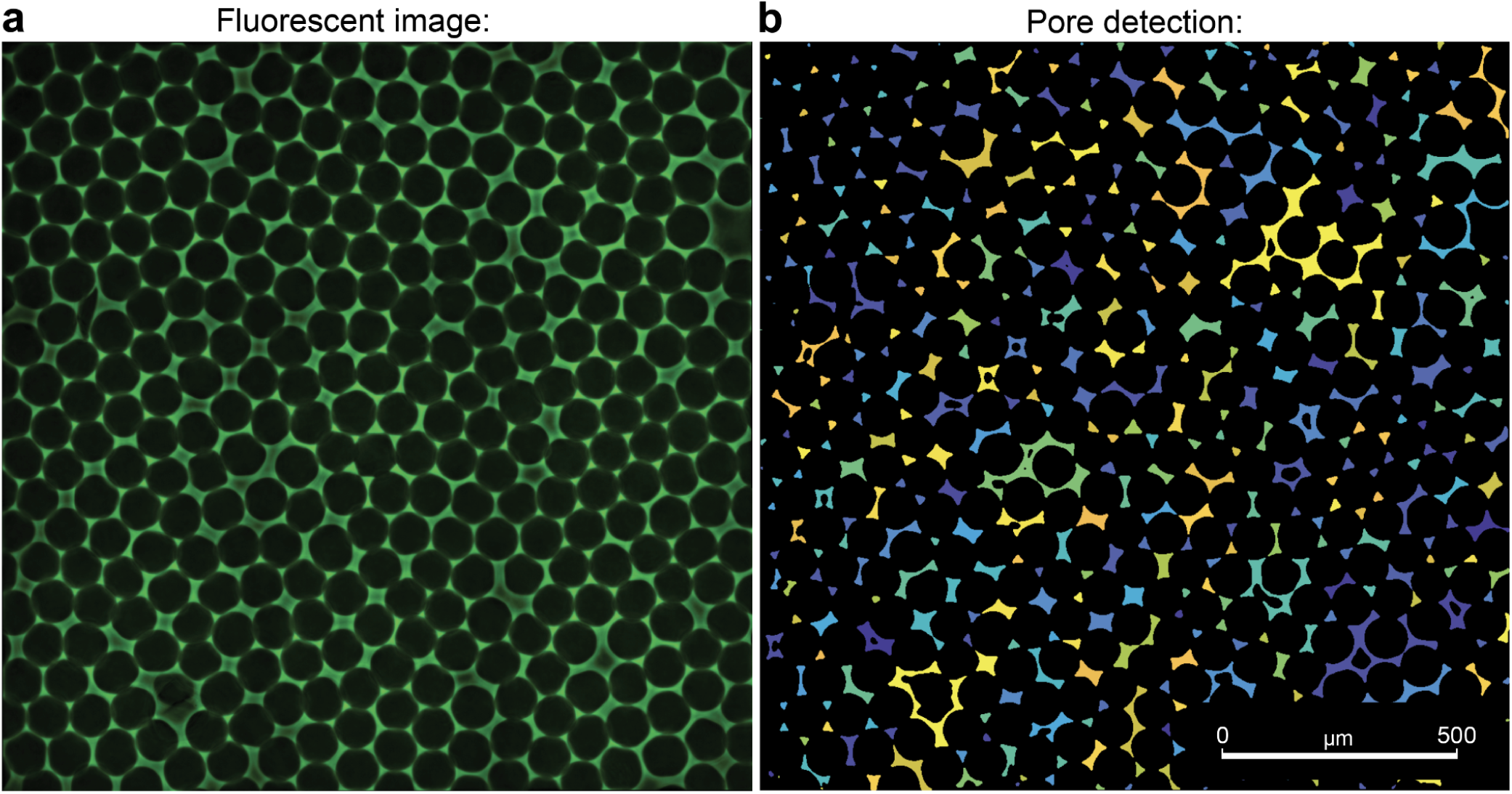
Equivalent median pore diameter analysis. **(a)** Transversal sections of GHS based on fluorescence microscopy. **(b)** Pore detection using a user-developed MATLAB code to identify the void spaces among photochemically assembled microgels occupied by fluorescently labeled dextran molecules (Mw = 2 MDa). To measure the equivalent pore diameter, detected pores were converted to circles with similar area, and the median of diameter distribution was calculated.

## Notes

### Competing Interest Statement

The authors have declared no competing interest.

## References

1. Paul, C. D., Mistriotis, P. & Konstantopoulos, K. Cancer cell motility: lessons from migration in confined spaces. Nat. Rev. Cancer 17, 131–140 (2017).

2. Provenzano, P. P. et al. Collagen reorganization at the tumor-stromal interface facilitates local invasion. BMC Med. 4, 38 (2006).

3. Rianna, C., Radmacher, M. & Kumar, S. Direct evidence that tumor cells soften when navigating confined spaces. MBoC 31, 1726–1734 (2020).

4. Balzer, E. M. et al. Physical confinement alters tumor cell adhesion and migration phenotypes. FASEB J. 26, 4045–4056 (2012).

5. Rolli, C. G., Seufferlein, T., Kemkemer, R. & Spatz, J. P. Impact of tumor cell cytoskeleton organization on invasiveness and migration: a microchannel-based approach. PLoS One 5, e8726 (2010).

6. Pathak, A. & Kumar, S. Independent regulation of tumor cell migration by matrix stiffness and confinement. Proc. Natl. Acad. Sci. U. S. A. 109, 10334–10339 (2012).

7. Raman, P. S., Paul, C. D., Stroka, K. M. & Konstantopoulos, K. Probing cell traction forces in confined microenvironments. Lab Chip 13, 4599–4607 (2013).

8. Liu, Y.-J. et al. Confinement and low adhesion induce fast amoeboid migration of slow mesenchymal cells. Cell 160, 659–672 (2015).

9. Ray, A., Slama, Z. M., Morford, R. K., Madden, S. A. & Provenzano, P. P. Enhanced Directional Migration of Cancer Stem Cells in 3D Aligned Collagen Matrices. Biophys. J. 112, 1023–1036 (2017).

10. Conklin, M. W. et al. Aligned collagen is a prognostic signature for survival in human breast carcinoma. Am. J. Pathol. 178, 1221–1232 (2011).

11. Provenzano, P. P. et al. Collagen density promotes mammary tumor initiation and progression. BMC Med. 6, 11 (2008).

12. van Zijl, F., Krupitza, G. & Mikulits, W. Initial steps of metastasis: Cell invasion and endothelial transmigration. Mutation Research/Reviews in Mutation Research 728, 23–34 (2011).

13. Tabdanov, E. D., Puram, V., Zhovmer, A. & Provenzano, P. P. Microtubule-Actomyosin Mechanical Cooperation during Contact Guidance Sensing. Cell Rep. 25, 328–338.e5 (2018).

14. Zhovmer, A. S. et al. Mechanical Counterbalance of Kinesin and Dynein Motors in a Microtubular Network Regulates Cell Mechanics, 3D Architecture, and Mechanosensing. ACS Nano (2021) doi:10.1021/acsnano.1c04435.

15. Tabdanov, E. D. et al. Bimodal sensing of guidance cues in mechanically distinct microenvironments. Nat. Commun. 9, 4891 (2018).

16. Kim, D., You, E. & Rhee, S. Dynein regulates cell migration depending on substrate rigidity. Int. J. Mol. Med. 29, 440–446 (2012).

17. Jayatilaka, H. et al. EB1 and cytoplasmic dynein mediate protrusion dynamics for efficient 3-dimensional cell migration. FASEB J. 32, 1207–1221 (2018).

18. Robison, P. et al. Detyrosinated microtubules buckle and bear load in contracting cardiomyocytes. Science 352, aaf0659 (2016).

19. Su, X., Li, H., Chen, S. & Qin, C. Study on the Prognostic Values of Dynactin Genes in Low-Grade Glioma. Technol. Cancer Res. Treat. 20, 15330338211010143 (2021).

20. Li, W. et al. Dynactin 2 acts as an oncogene in hepatocellular carcinoma through promoting cell cycle progression. Liver Research 6, 155–166 (2022).

21. Bransfield, K. L., Askham, J. M., Leek, J. P., Robinson, P. A. & Mighell, A. J. Phenotypic changes associated with DYNACTIN-2 (DCTN2) over expression characterise SJSA-1 osteosarcoma cells. Mol. Carcinog. 45, 157–163 (2006).

22. Wang, Q. et al. Prognostic Value of Dynactin mRNA Expression in Cutaneous Melanoma. Med. Sci. Monit. 24, 3752–3763 (2018).

23. Gong, L.-B. et al. DYNC1I1 Promotes the Proliferation and Migration of Gastric Cancer by Up-Regulating IL-6 Expression. Front. Oncol. 9, 491 (2019).

24. Hassan Ibrahim, I., Balah, A., Gomaa Abd Elfattah Hassan, A. & Gamal Abd El-Aziz, H. Role of motor proteins in human cancers. Saudi J. Biol. Sci. 29, 103436 (2022).

25. Uhlen, M. et al. A pathology atlas of the human cancer transcriptome. Science 357, (2017).

26. Koster, J., Volckmann, R., Zwijnenburg, D., Molenaar, P. & Versteeg, R. R2: Genomics analysis and visualization platform. R2: Genomics Analysis and Visualization Platform (http://r2.amc.nl http://r2platform.com).

27. Doyle, A. D., Wang, F. W., Matsumoto, K. & Yamada, K. M. One-dimensional topography underlies three-dimensional fibrillar cell migration. J. Cell Biol. 184, 481–490 (2009).

28. Guetta-Terrier, C. et al. Protrusive waves guide 3D cell migration along nanofibers. J. Cell Biol. 211, 683–701 (2015).

29. Ray, A. et al. Anisotropic forces from spatially constrained focal adhesions mediate contact guidance directed cell migration. Nat. Commun. 8, 14923 (2017).

30. Han, W. et al. Oriented collagen fibers direct tumor cell intravasation. Proceedings of the National Academy of Sciences 113, 11208–11213 (2016).

31. Wang, W. Y., Davidson, C. D., Lin, D. & Baker, B. M. Actomyosin contractility-dependent matrix stretch and recoil induces rapid cell migration. Nat. Commun. 10, 1–12 (2019).

32. Wang, W. Y. et al. Extracellular matrix alignment dictates the organization of focal adhesions and directs uniaxial cell migration. APL Bioeng 2, 046107 (2018).

33. Geiger, F., Rüdiger, D., Zahler, S. & Engelke, H. Fiber stiffness, pore size and adhesion control migratory phenotype of MDA-MB-231 cells in collagen gels. PLoS One 14, e0225215 (2019).

34. Doyle, A. D., Sykora, D. J., Pacheco, G. G., Kutys, M. L. & Yamada, K. M. 3D mesenchymal cell migration is driven by anterior cellular contraction that generates an extracellular matrix prestrain. Dev. Cell 56, 826–841.e4 (2021).

35. Yamada, K. M. & Sixt, M. Mechanisms of 3D cell migration. Nat. Rev. Mol. Cell Biol. 20, 738–752 (2019).

36. Cai, Y. et al. Cytoskeletal coherence requires myosin-IIA contractility. J. Cell Sci. 123, 413–423 (2010).

37. Lomakin, A. J. et al. Competition for actin between two distinct F-actin networks defines a bistable switch for cell polarization. Nat. Cell Biol. 17, 1435–1445 (2015).

38. Liu, Z. et al. Blebbistatin inhibits contraction and accelerates migration in mouse hepatic stellate cells. Br. J. Pharmacol. 159, 304–315 (2010).

39. Doyle, A. D. et al. Micro-environmental control of cell migration--myosin IIA is required for efficient migration in fibrillar environments through control of cell adhesion dynamics. J. Cell Sci. 125, 2244–2256 (2012).

40. Kastelic, J., Galeski, A. & Baer, E. The Multicomposite Structure of Tendon. Connect. Tissue Res. 6, 11–23 (1978).

41. Ramirez-San Juan, G. R., Oakes, P. W. & Gardel, M.L. Contact guidance requires spatial control of leading-edge protrusion. Mol. Biol. Cell 28, 1043–1053 (2017).

42. Höing, S. et al. Dynarrestin, a Novel Inhibitor of Cytoplasmic Dynein. Cell Chem Biol 25, 357–369.e6 (2018).

43. Murrell, M., Oakes, P. W., Lenz, M. & Gardel, M. L. Forcing cells into shape: the mechanics of actomyosin contractility. Nat. Rev. Mol. Cell Biol. 16, 486–498 (2015).

44. Kovács, M., Tóth, J., Hetényi, C., Málnási-Csizmadia, A. & Sellers, J. R. Mechanism of blebbistatin inhibition of myosin II. J. Biol. Chem. 279, 35557–35563 (2004).

45. Roman, B. I., Verhasselt, S. & Stevens, C. V. Medicinal Chemistry and Use of Myosin II Inhibitor (S)-Blebbistatin and Its Derivatives. J. Med. Chem. 61, 9410–9428 (2018).

46. Roossien, D. H., Miller, K. E. & Gallo, G. Ciliobrevins as tools for studying dynein motor function. Front. Cell. Neurosci. 9, 252 (2015).

47. Steinman, J. B. et al. Chemical structure-guided design of dynapyrazoles, cell-permeable dynein inhibitors with a unique mode of action. Elife 6, (2017).

48. Kerr, J. P. et al. Detyrosinated microtubules modulate mechanotransduction in heart and skeletal muscle. Nat. Commun. 6, 1–14 (2015).

49. Jimenez, A. J. et al. Acto-myosin network geometry defines centrosome position. Curr. Biol. (2021) doi:10.1016/j.cub.2021.01.002.

50. Farina, F. et al. The centrosome is an actin-organizing centre. Nat. Cell Biol. 18, 65–75 (2016).

51. Fokin, A. I. et al. The Arp1/11 minifilament of dynactin primes the endosomal Arp2/3 complex. Sci Adv 7, (2021).

52. Pimm, M. L. & Henty-Ridilla, J. L. New twists in actin-microtubule interactions. Mol. Biol. Cell 32, 211–217 (2021).

53. Kubow, K. E., Shuklis, V. D., Sales, D. J. & Horwitz, A. R. Contact guidance persists under myosin inhibition due to the local alignment of adhesions and individual protrusions. Sci. Rep. 7, 14380 (2017).

54. Randall, T. S. et al. A small-molecule activator of kinesin-1 drives remodeling of the microtubule network. Proc. Natl. Acad. Sci. U. S. A. 114, 13738–13743 (2017).

55. Cailleau, R., Olivé, M. & Cruciger, Q. V. Long-term human breast carcinoma cell lines of metastatic origin: preliminary characterization. In Vitro 14, 911–915 (1978).

56. Kalluri, R. & Weinberg, R. A. The basics of epithelial-mesenchymal transition. J. Clin. Invest. 119, 1420–1428 (2009).

57. Lee, A. V., Oesterreich, S. & Davidson, N. E. MCF-7 cells--changing the course of breast cancer research and care for 45 years. J. Natl. Cancer Inst. 107, (2015).

58. Nohara, K., Wang, F. & Spiegel, S. Glycosphingolipid composition of MDA-MB-231 and MCF-7 human breast cancer cell lines. Breast Cancer Res. Treat. 48, 149–157 (1998).

59. Uhlén, M. et al. Tissue-based map of the human proteome. Science 347, 1260419 (2015).

60. Rhee, S., Jiang, H., Ho, C.-H. & Grinnell, F. Microtubule function in fibroblast spreading is modulated according to the tension state of cell--matrix interactions. Proceedings of the National Academy of Sciences 104, 5425–5430 (2007).

61. Bershadsky, A., Kozlov, M. & Geiger, B. Adhesion-mediated mechanosensitivity: a time to experiment, and a time to theorize. Curr. Opin. Cell Biol. 18, 472–481 (2006).

62. McGrail, D. J., Mezencev, R., Kieu, Q. M. N., McDonald, J. F. & Dawson, M. R. SNAIL-induced epithelial-to-mesenchymal transition produces concerted biophysical changes from altered cytoskeletal gene expression. FASEB J. 29, 1280–1289 (2015).

63. Gracia, M. et al. Mechanical impact of epithelial−mesenchymal transition on epithelial morphogenesis in Drosophila. Nat. Commun. 10, 1–17 (2019).

64. Riching, K. M. et al. 3D collagen alignment limits protrusions to enhance breast cancer cell persistence. Biophys. J. 107, 2546–2558 (2014).

65. del Castillo, U., Winding, M., Lu, W. & Gelfand, V. I. Interplay between kinesin-1 and cortical dynein during axonal outgrowth and microtubule organization in Drosophila neurons. Elife 4, e10140 (2015).

66. Shakiba, D. et al. The Balance between Actomyosin Contractility and Microtubule Polymerization Regulates Hierarchical Protrusions That Govern Efficient Fibroblast–Collagen Interactions. ACS Nano 14, 7868–7879 (2020).

67. Grabham, P. W., Seale, G. E., Bennecib, M., Goldberg, D. J. & Vallee, R. B. Cytoplasmic dynein and LIS1 are required for microtubule advance during growth cone remodeling and fast axonal outgrowth. J. Neurosci. 27, 5823–5834 (2007).

68. Laan, L. et al. Cortical Dynein Controls Microtubule Dynamics to Generate Pulling Forces that Position Microtubule Asters. Cell 148, 502–514 (2012).

69. Monzon, G. A. et al. Stable tug-of-war between kinesin-1 and cytoplasmic dynein upon different ATP and roadblock concentrations. J. Cell Sci. 133, (2020).

70. Roberts, A. J., Kon, T., Knight, P. J., Sutoh, K. & Burgess, S. A. Functions and mechanics of dynein motor proteins. Nat. Rev. Mol. Cell Biol. 14, 713–726 (2013).

71. Adio, S., Reth, J., Bathe, F. & Woehlke, G. Review: regulation mechanisms of Kinesin-1. J. Muscle Res. Cell Motil. 27, 153–160 (2006).

72. Ataie, Z. et al. Nanoengineered Granular Hydrogel Bioinks with Preserved Interconnected Microporosity for Extrusion Bioprinting. Small 18, e2202390 (2022).

73. Ataie, Z., Jaberi, A., Kheirabadi, S., Risbud, A. & Sheikhi, A. Gelatin Methacryloyl Granular Hydrogel Scaffolds: High-throughput Microgel Fabrication, Lyophilization, Chemical Assembly, and 3D Bioprinting. J. Vis. Exp. (2022) doi:10.3791/64829.

74. Sheikhi, A. et al. Microfluidic-enabled bottom-up hydrogels from annealable naturally-derived protein microbeads. Biomaterials 192, 560–568 (2019).

75. Yang, D. et al. Promoting cell migration in tissue engineering scaffolds with graded channels. Adv. Healthc. Mater. 6, 1700472 (2017).

76. Prager-Khoutorsky, M., Khoutorsky, A. & Bourque, C. W. Unique interweaved microtubule scaffold mediates osmosensory transduction via physical interaction with TRPV1. Neuron 83, 866–878 (2014).

77. Tabdanov, E. D., Zhovmer, A. S., Puram, V. & Provenzano, P. P. Engineering Elastic Nano-and Micro-Patterns and Textures for Directed Cell Motility. STAR Protocols (2020).

78. Schmid, H. & Michel, B. Siloxane Polymers for High-Resolution, High-Accuracy Soft Lithography. Macromolecules 33, 3042–3049 (2000).

79. Plotnikov, S. V., Sabass, B., Schwarz, U. S. & Waterman, C. M. High-resolution traction force microscopy. Methods Cell Biol. 123, 367–394 (2014).

80. Odom, T. W., Love, J. C., Wolfe, D. B., Paul, K. E. & Whitesides, G. M. Improved Pattern Transfer in Soft Lithography Using Composite Stamps. Langmuir 18, 5314–5320 (2002).

81. Fischer, R. S., Myers, K. A., Gardel, M. L. & Waterman, C. M. Stiffness-controlled three-dimensional extracellular matrices for high-resolution imaging of cell behavior. Nat. Protoc. 7, 2056–2066 (2012).

82. Plotnikov, S. V., Sabass, B., Schwarz, U. S. & Waterman, C. M. High-resolution traction force microscopy. Methods Cell Biol. 123, 367–394 (2014).

83. Schindelin, J. et al. Fiji: an open-source platform for biological-image analysis. Nat. Methods 9, 676–682 (2012).

